# Annealing synchronizes the 70*S* ribosome into a minimum-energy conformation

**DOI:** 10.1101/2021.06.25.447849

**Authors:** Xiaofeng Chu, Xin Su, Mingdong Liu, Li Li, Tianhao Li, Yicheng Qin, Guoliang Lu, Lei Qi, Yunhui Liu, Jinzhong Lin, Qing-Tao Shen

## Abstract

Researchers commonly anneal metals, alloys, and semiconductors to repair defects and improve microstructures via recrystallization. Theoretical studies indicate simulated annealing on biological macromolecules helps predict the final structures with minimum free energy. Experimental validation of this homogenizing effect and further exploration of its applications are fascinating scientific questions that remain elusive. Here, we chose the apo-state 70*S* ribosome from *Escherichia coli* as a model, wherein the 30*S* subunit undergoes a thermally driven inter-subunit rotation and exhibits substantial structural flexibility as well as distinct free energy. We experimentally demonstrate that annealing at a fast cooling rate enhances the 70*S* ribosome homogeneity and improves local resolution on the 30*S* subunit. After annealing, the 70*S* ribosome is in a nonrotated state with respect to corresponding intermediate structures in unannealed or heated ribosomes, and exhibits a minimum energy in the free energy landscape. One can readily crystallize these minimum-energy ribosomes, which have great potential for synchronizing proteins on a single-molecule level. Our experimental results are consistent with theoretical analysis on the temperature-dependent Boltzmann distribution, and offer a facile yet robust approach to enhance protein stability, which is ideal for high-resolution cryogenic electron microscopy. Beyond structure determination, annealing can be extended to study protein folding and explore conformational and energy landscape.

**Significance statement:** In metallurgy, annealing heats a metal or alloy to a predetermined temperature, holding for a certain time, and then cooling to room temperature to change the physical and sometimes also the chemical properties of the material. Researchers introduce the similar concept as simulated annealing to predict minimum-energy conformations of biological macromolecules. In this work, we experimentally verify that annealing at a fast cooling rate can synchronize the 70*S* ribosome into a nonrotated state with a minimum energy in the free energy landscape. Our results not only offer a facile yet robust approach to stabilize proteins for high-resolution structural analysis, but also contribute to the understanding of protein folding and temperature adaptation.

## INTRODUCTION

Annealing—common in metallurgy—heats a metal or alloy to a set temperature, holds this temperature, and then cools to room temperature to improve the physical and sometimes also the chemical properties of the material (1, 2). During annealing, recrystallization repairs defects to afford a refined microstructure, which can relieve stress, soften the metal, increase the ductility, and improve the grain structure. One can also anneal organic/inorganic semiconductors via rearranging molecules, polymer chain segments, or entire polymer chains into active layer films. Optimizing the annealing temperature and time can either dramatically increase the crystal size from the sub-micrometer scale to several micrometers, or yield different crystals (3–5).

Annealed materials tend to adopt homogenous states and readily assemble into either three-dimensional (3D) or two-dimensional (2D) crystals. One can readily visualize such regular packing via atomic force microscopy (AFM), X-ray diffraction (XRD), or electron microscopy (EM). Whether annealing exhibits similar effects on biological macromolecules, especially proteins, is a fascinating scientific question that remains unanswered. To date, annealing on proteins has mainly been limited to molecular dynamics simulations. Researchers have simulated annealing—such as with Monte Carlo algorithms—on proteins from a set of starting conformations, which has helped predict the final structures with minimum free energy (6–8). Distinct from metals and organic polymers, proteins and protein complexes are usually discrete entities that consist of chemically diverse subunits bound together in diverse geometries. This substantial structural heterogeneity hinders direct structural determination via AFM or XRD. In contrast, recent advancements in the resolution of cryo-EM offer great opportunities to obtain high-resolution protein structures at the single-molecule level (9–12). By using cryo-EM to compare detailed structures before/after annealing, one can obtain direct experimental evidence for annealing to affect protein conformations.

Many proteins, especially protein complexes, can in principle exhibit diverse conformations. Based on the Boltzmann distribution formula, each conformational state of a protein or protein complex is occupied with a statistical weight, *P_i_* = *e^-^*^(*E_i_*/*k_B_T*)^, where *E_i_* is the free energy of the conformational state *i*, *k_B_* is the Boltzmann constant, and *T* is the absolute temperature. The Boltzmann distribution and the derived free energy are temperature-dependent; an increase in temperature can directly increase the free energy and help a protein escape the energy barrier between local minima. At room and physiological temperatures, minimum-energy conformations will usually differ from those at lower temperatures. This is the theoretical basis for protein conformational changes in accordance with temperature.

## RESULTS

### Annealing improves local resolution

Here, we chose the apo-state 70*S* ribosome from *Escherichia coli* as a model, wherein the 30*S* subunit undergoes a thermally driven inter-subunit rotation (13–18) and exhibits substantial structural flexibility as well as distinct free energy (19). We incubated purified apo-state 70*S* ribosome at 0°C for 5 min, then immediately flash-froze the ribosome for cryo-EM analysis, which retained the same conformation as before vitrification (depicted as the unannealed state). We screened the collected 70*S* ribosome particles via discarding obvious junk and disassembled ribosomes through 2D and 3D classifications. Reconstruction from 200,000 randomly selected particles yielded a structure at a final resolution of 2.6 Å, based on gold-standard Fourier shell correlation (FSC; Fig. 1A and figs. S1-3). Due to the lack of stabilizing factors, such as messenger and transfer RNA, local resolution estimates on the unannealed 70*S* ribosome indicated variable resolutions across the entire density map over the 2.6–7.2 Å range (Fig. 1A). Relative to the 50*S* subunit, the 30*S* subunit—especially its head domain—was less well-resolved, which is common in other apo-state ribosomes (20, 21).

**Fig. 1.**
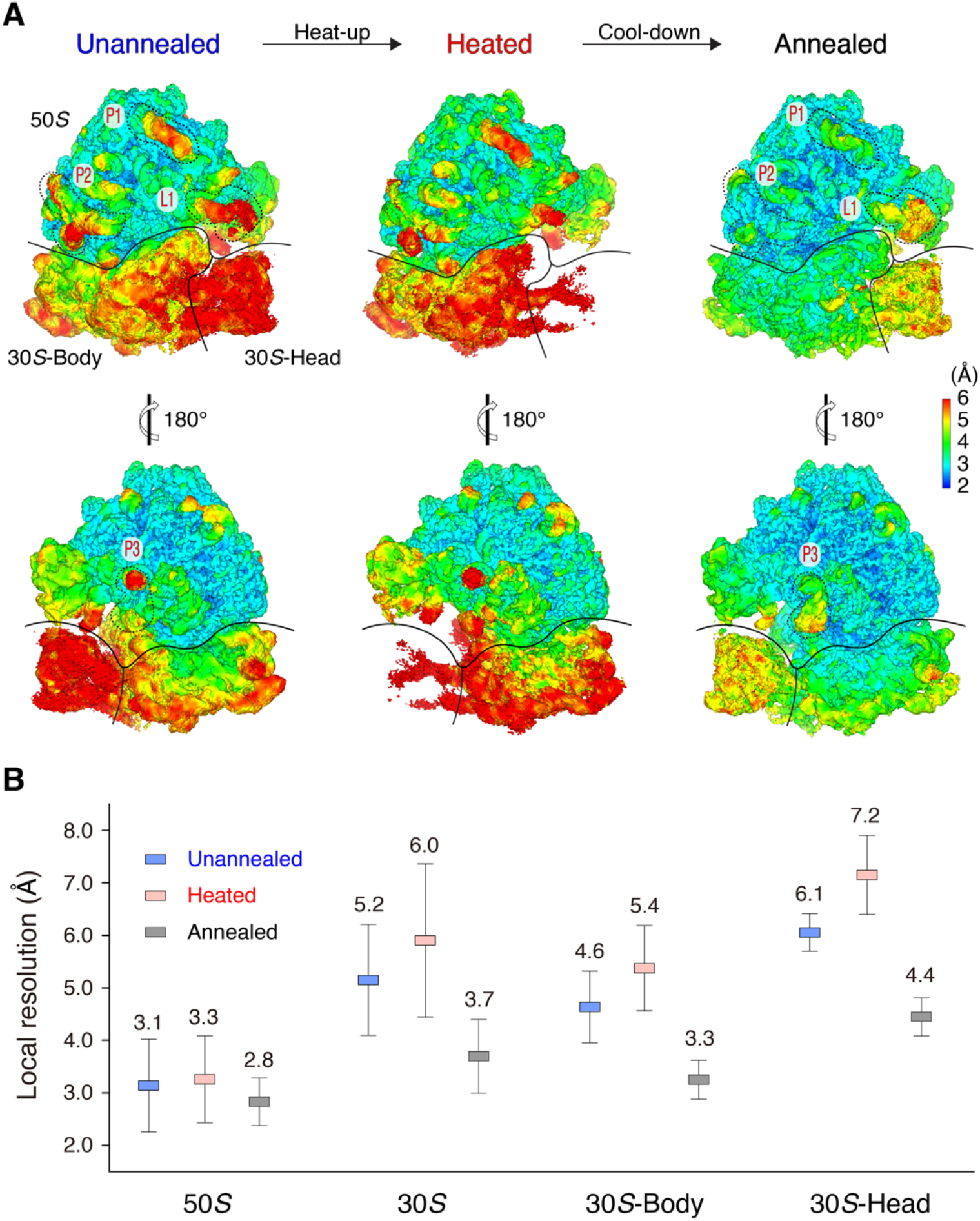
Annealing improves local resolution of the 70*S* ribosome. (**A**) Local resolution maps of the unannealed, heated, and annealed 70*S* ribosomes. The 50*S* subunit and the body and head domains of the 30*S* subunit are labeled. L1 stalk (L1) and other three sub-regions (P1, P2, and P3) on the periphery of the 50*S* subunit in the unannealed and annealed 70*S* ribosomes are marked for resolution comparison. (**B**) Local resolution comparison of the unannealed, heated, and annealed 70*S* ribosomes.

To quantitate the resolution variation of different regions, we calculated the local resolution via averaging local resolution values within the selected regions (figs. S4-6). Our analysis indicated that the 50*S* subunit exhibited an average local resolution of 3.1 Å, whereas the 30*S* subunit was much less resolved—only 5.2 Å. Furthermore, the 30*S* head domain was even less resolved—average resolution of 6.1 Å (Fig. 1B). The inter-subunit ratcheting between the 50*S* and 30*S* subunits is the primary cause for poor resolvability; the intra-subunit swirling of the 30*S* subunit is secondary, which lessens the resolution of the head domain. For simplicity, we used the local resolution of the 30*S* subunit as a marker to monitor the effect of annealing on the 70*S* ribosome.

Full annealing consists of two tandem heating and cooling steps. Cooling rate and annealing temperature are usually two critical elements for homogenization of metals, alloys, and semiconductors (22, 23). In metallurgy, the annealing temperature is usually 30°C to 50°C greater than the upper critical temperature of metals (24). In molecular dynamics simulation, researchers sometimes use extreme annealing temperatures—such as 127°C—to help biological macromolecules escape energy barriers. To avoid heat-induced denaturation, we set the annealing temperatures of the 70*S* ribosome below the ∼72°C melting temperature of bacterial ribosomes (25, 26). We carried out subsequent cooling on heated ribosomes in highly thermo-conductive PCR tubes, via either gradual cooling in the PCR machine or immediate immersion into an ice bath. We vitrified the annealed samples in a Vitrobot chamber preset at 4°C, and then subjected the samples to single-particle cryo-EM analysis in accordance with the same procedure used for unannealed ribosomes from the exact same number (200, 000) of randomly selected particles (Fig. 2A).

**Fig. 2.**
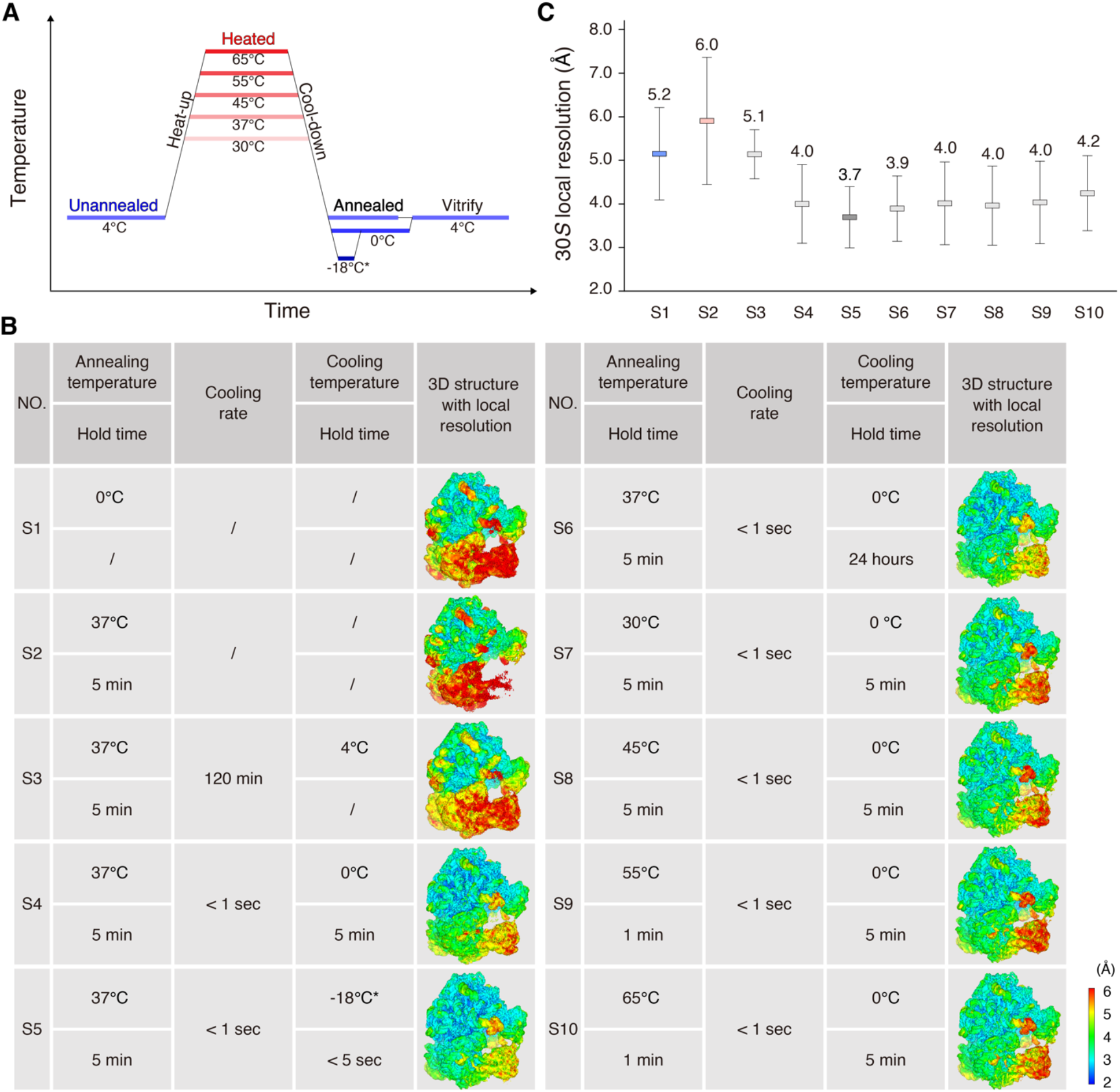
Systematic screening on annealing conditions. (**A**) Schematic for annealing of the 70*S* ribosome. The unannealed, heated, and annealed stages are labeled on the diagram. (**B**) Detailed annealing conditions on the 70*S* ribosomes and the respective 3D local resolution maps. We used sample names S1–S10 in (**C**) and other figures. Regarding S5, we briefly (<5 s) immersed the 70*S* ribosome into a mixture of salt, ice, and water pre-cooled to −18°C, and then transferred the samples to a mixture of ice and water at 0°C before freezing. (**C**) Local resolution comparison of 30*S* subunits among various annealing conditions.

We performed systematic screening of various combinations of cooling rates and annealing temperatures via comparing the local resolutions of the 30*S* subunit. When we gradually cooled the 70*S* ribosome over the course of 2 h from 37°C to 0°C, the average local resolution of the 30*S* subunit was 5.1 Å, and there was no obvious local resolution improvement with respect to the unannealed ribosome—5.2 Å (Fig. 2B, C and fig. S6). Fast cooling via immersion into an ice bath (<1 s) improved the local resolution from a value of 5.2 Å to a value of ∼4.0 Å on the 30*S* subunit (Fig. 2B, C and fig. S6). A mixture of salt, ice, and water— comparatively high thermo-conductivity—yielded improved local resolution of the 30*S* subunit—3.7 Å; the 30*S* head was also dramatically improved to 4.4 Å, relative to 6.1 Å from the unannealed ribosome (Figs. 1B and 2B, C and fig. S6). Apparently, one can use fast instead of gradual cooling to improve the local resolution of the 30*S* subunit. As another critical parameter for annealing in metallurgy, annealing temperatures of 45°C, 55°C, and 65°C did not further improve local resolutions of the 30*S* subunit, compared with 30°C and 37°C (Fig. 2B, C and fig. S6). Thus, we considered fast cooling from 37°C in a mixture of salt, ice, and water as ideal annealing conditions for improving the local resolution of the 70*S* ribosome.

To further understand the annealing process, we also investigated heating the 70*S* ribosome to an annealing temperature of 37°C as an intermediate state by cryo-EM (Fig. 2A). Researchers recently developed temperature-resolved cryo-EM to study temperature-sensitive proteins, such as transient receptor potential vanilloid-3 (TRPV3) and ketol-acid reductoisomerase (KARI), at optimal temperatures (27, 28). We optimized sample preparation, such as by consistent high-temperature control and efficient prevention of ice contamination, and found these procedures useful for obtaining ideal cryo-EM grids (fig. S7 and movie S1). We resolved 200,000 particles of the heated 70*S* ribosome at a resolution of 2.7 Å, close to that of the unannealed ribosome. However, the average local resolution of the 30*S* subunit was 6.0 Å, substantially less than that of the unannealed and annealed structures (Figs. 1B and 2B, C and fig. S6). The 30*S* head domain was only resolved at 7.2 Å, and the densities were almost indiscernible at the threshold level proper to the 50*S* subunit (Figs. 1 and 2B, C and fig. S6). Thus, heating increased the free energy of the 70*S* ribosome and imparted more structural flexibility to the 30*S* subunit, which to some extent hindered final reconstruction and afforded inferior resolution.

### Annealing stabilizes flexible regions

Relative to the unannealed 70*S* ribosome, the overall structure of the annealed 70*S* ribosome exhibited substantially higher resolution. The resolution improvement was not uniform across the entire 70*S* ribosome. For instance, the 30*S* subunit, which was poorly resolved in the unannealed ribosome, was dramatically improved 1.5 Å—i.e., from a value of 5.2 Å to a value of 3.7 Å. As the control, the well-resolved 50*S* subunit was only enhanced 0.3 Å—i.e., from a value of 3.1 Å to a value of 2.8 Å—after annealing (Fig. 1B). Thus, annealing was especially beneficial to low-resolution regions, which have greater structural flexibility. To further verify this deduction, we performed a comprehensive statistical analysis on the average local resolution of the same sub-regions between the unannealed and annealed 70*S* ribosomes. For example, annealing improved the average local resolutions of different regions at the level of ∼0.1, 0.6, 0.8, 1.2, and 2.0 Å; the corresponding regions in the unannealed ribosome had local resolution in the range of 2.5–3.0, 3.0–3.5, 4.0–4.5, 5.0–5.5, and 5.5–6.0 Å (Fig. 3A). An exponential curve fit well to the data, indicating that more flexibility in the unannealed 70*S* ribosome corresponded to more improvement in local resolution after annealing.

**Fig. 3.**
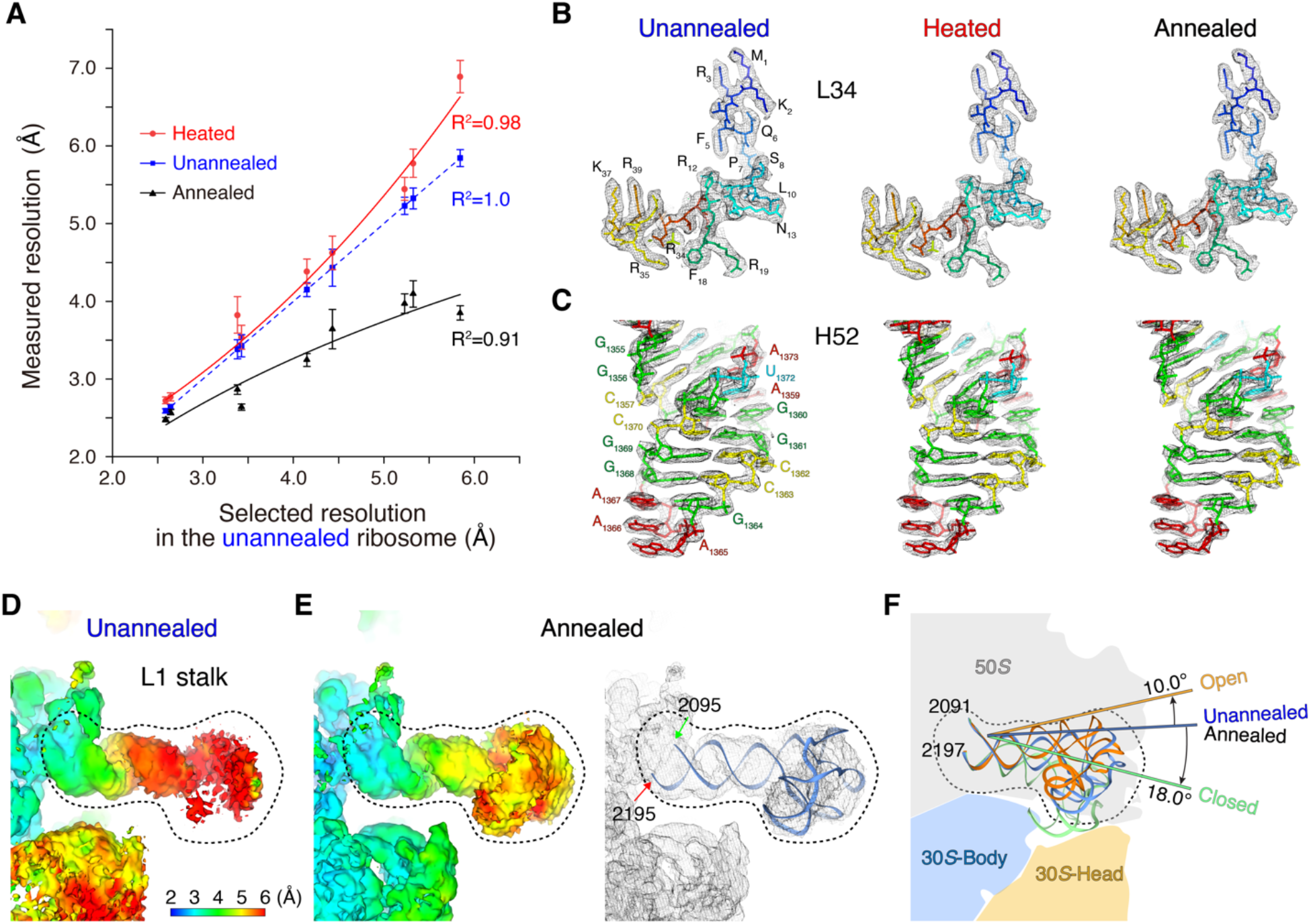
Annealing stabilizes flexible regions of the 70*S* ribosome. (**A**) The effects of heating and annealing on regions with different local resolutions. We selected regions with different local resolutions from 2.5–6.0 Å in the unannealed 70*S* ribosome, and measured local resolutions in the corresponding regions in the heated and annealed 70*S* ribosomes. (**B**) and (**C**) Cryo-EM maps and fitted atomic models for L34 ribosomal protein and H52 helix in 23*S* rRNA under unannealed, heated, and annealed conditions. (**D**) Local resolution map of the L1 stalk in the unannealed ribosome. The position of the L1 stalk is shown in Fig. 1A. (**E**) Local resolution map of the L1 stalk in the annealed ribosome and fitted atomic model. (**F**) Intermediate states of the L1 stalk in the unannealed and annealed ribosomes, with respect to the known open and closed states. Atomic models of the L1 stalk are as follows: open (gold), intermediate (blue), and closed (green) states.

Rigid regions were well-resolved in both unannealed and annealed 70*S* ribosomes, and most RNA fragments of 23*S* rRNA and ribosomal proteins in the 50*S* subunit were clearly resolved in the cryo-EM maps (Fig. 3B, C). There were some flexible peripheral regions in the 50*S* subunit, which were poorly resolved in the unannealed ribosome. These regions were substantially more distinct after annealing (Fig. 1A). For instance, the L1 stalk, a mobile domain of the 50*S* subunit that shields the exit site of the deacylated tRNA, is usually invisible in X-ray crystallography and cryo-EM, especially in the absence of peptidyl–tRNA and aminoacyl–tRNA (29–31). After annealing, the L1 stalk was resolved at a resolution of 4.9 Å, and adopted an orientation between the open and closed states, which researchers classify as an intermediate state during translation elongation (Fig. 3D-F). In addition to the L1 stalk, regions from 1,034– 1,121 and 1,437–1,553 in 23*S* rRNA are intrinsically flexible, such that they interact with EF–G– GTP (during elongation and termination procedures) and EF–Tu–GTP–aminoacyl–tRNA (help coupling with the SecY complex). The resolution of these flexible regions was substantially improved: from a value of 4.9 Å to a value of 4.1 Å, and from a value of 4.0 Å to a value of 2.9 Å, respectively, after annealing (fig. S8).

Another resolution improvement after annealing was evident on the flexible 30*S* subunit. In the annealed ribosome, the 30*S* body domain was improved 1.3 Å: i.e., from a value of 4.6 Å to a value of 3.3 Å (Figs. 1B and 2B). Furthermore, swirling of the 30*S* head domain, with respect to the 30*S* body domain, imparted more structural flexibility to the 30*S* head domain; we resolved the unannealed 30*S* head domain to only 6.1 Å. After annealing, we dramatically stabilized the 30*S* head domain and improved the resolution to 4.4 Å (Figs. 1B and 2B). The resolution improvement in the 30*S* subunit was also in accordance with our previous finding that greater structural flexibility corresponded to greater resolution improvement after annealing.

Compared with the unannealed ribosome, the overall structure of the heated ribosome was less well-resolved. The resolution loss also occurred mainly on the flexible regions of the 70*S* ribosome. Similar local resolution comparisons between the unannealed and heated ribosomes indicated that more flexibility corresponded to a greater extent of resolution loss after heating. Briefly, the stable 50*S* subunit in the heated 70*S* ribosome had a similar resolution, at ∼3.3 Å, to the unannealed ribosome; whereas the flexible 30*S* subunit was substantially less well-resolved, from 5.2–6.0 Å (Figs. 1B and 2B and fig. S8). Apparently, heating had a more substantial influence on the flexible rather than rigid regions, analogous to the effect of annealing. At low-resolution ranges, such as 5–6 Å, in the unannealed ribosome, the resolution improvement via heating was ∼1 Å. This was not as large as the ∼2-Å improvement upon annealing, which indicates that protein structure may be more sensitive to annealing than heating (Fig. 3A).

### Annealing renders a different state

In metallurgy, recrystallization repairs defects and refines the microstructures after annealing. Thus, we tested whether the annealed 70*S* ribosome underwent conformational changes. To sample such changes before/after annealing, one requires accurate modeling— especially on the 30*S* subunit. We separately fit the 50*S* and 30*S* subunits from the crystallographic model (PDB ID 5MDZ) into the unannealed, heated, and annealed cryo-EM density maps via rigid body docking. Even though the fitting of the 30*S* head domain into the unannealed and heated 70*S* ribosomes was not as ideal as that of the annealed 70*S* ribosome, the well-resolved 30*S* body domains ensured docking accuracy among the unannealed, heated, and annealed 70*S* ribosomes (fig. S9).

We aligned the unannealed, heated, and annealed 70*S* ribosomes using the 50*S* subunit as a reference. We compared the orientations of 30*S* subunits via calculating the rotation angles relative to each other. The annealed 30*S* subunit rotated clockwise 7.4°—relative to the unannealed 30*S* subunit—exhibiting a distinct orientation. The motions of the 30*S* subunit after annealing resulted in shifts at the periphery of the ribosome at 12.0 Å (Fig. 4A). Annealing imparted a new orientation to the 70*S* ribosome and rendered the 30*S* subunit into a new state. The orientations of the 30*S* subunits between the unannealed and heated 70*S* ribosomes were close, with a variation less than 1.8 Å (Fig. 4B). This further indicates that the annealing-induced orientation change occurs during cooling instead of heating.

**Fig. 4.**
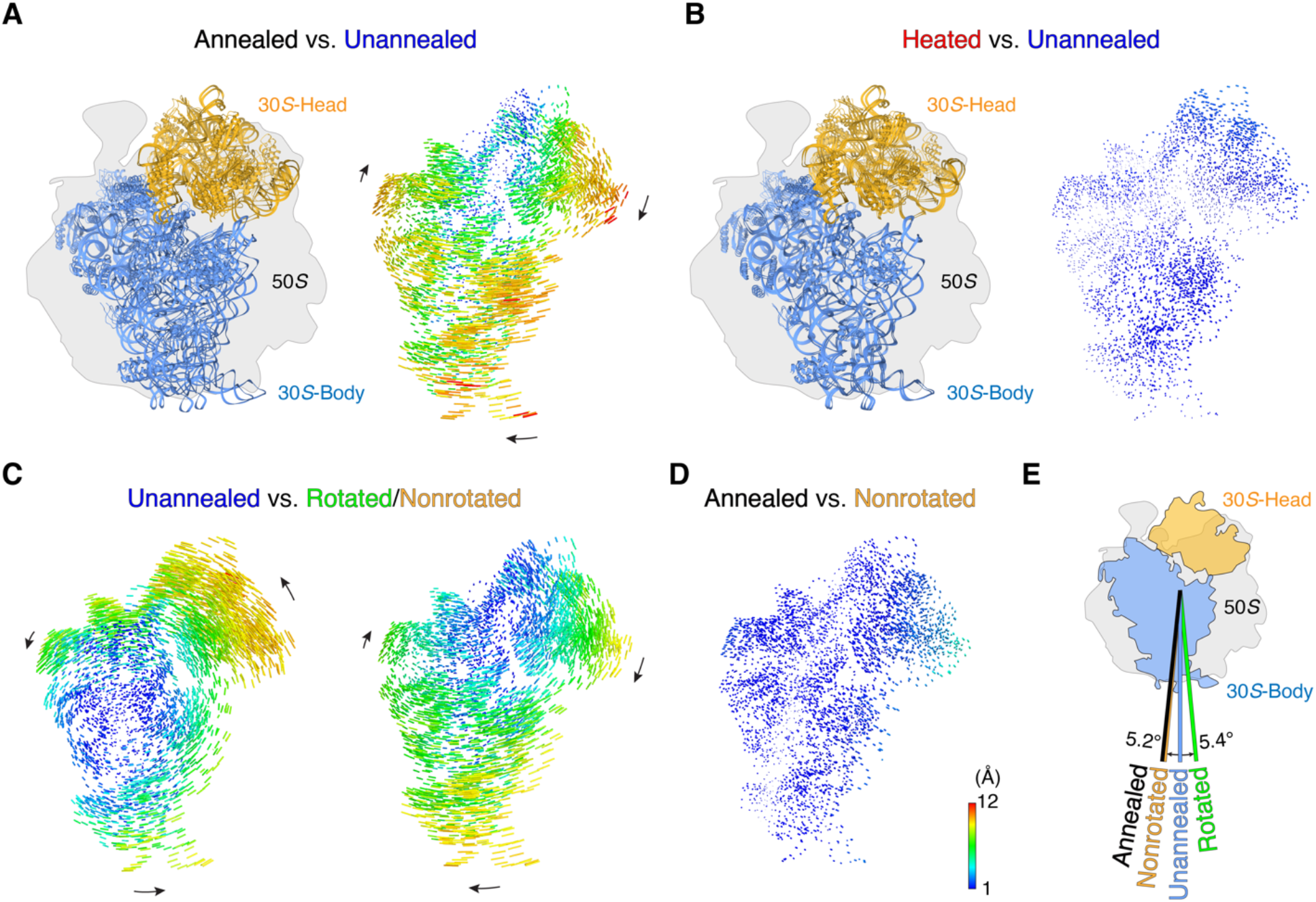
Annealing renders the 70*S* ribosome into a nonrotated state. (**A**) Rotational comparison of 30*S* subunits between the annealed and unannealed ribosomes. Left: We aligned the 50*S* subunits in the annealed and unannealed ribosomes as a reference, and atomic models for the 30*S* subunits are shown. Right: Difference vectors between phosphorous and C*α* atoms in the 30*S* subunits, with arrows indicating the direction of the change. (**B**) Rotational comparison of 30*S* subunits between the heated and unannealed ribosomes. (**C**) Rotational comparison of 30*S* subunit in the unannealed ribosome against the known fully rotated (PDB ID 4V9D, Left) and nonrotated (PDB ID 4V7C, Right) states. (**D**) Rotational comparison of 30*S* subunit in the annealed ribosome against the nonrotated state. (**E**) Summary of rotation angles of 30*S* subunit among the unannealed and annealed ribosomes, and the rotated and nonrotated states.

Ratcheting of the 30*S* subunit relative to the 50*S* subunit is the major rotation that facilitates tRNA translocation. Researchers thoroughly investigated the nonrotated (PDB ID 4V9D) and fully rotated (PDB ID 4V7C) states of the 30*S* subunit in the context of nascent peptide synthesis (32, 33). We aligned the unannealed, heated, and annealed ribosomes with both the nonrotated and fully rotated states, using the 50*S* subunits as a reference. In the absence of mRNA, tRNA, and other factors, the unannealed 30*S* subunit adopted an intermediate state between the nonrotated and fully rotated states. Specifically, the unannealed 30*S* subunit reached the fully rotated state via a 5.4° anticlockwise rotation, and clockwise rotated 5.2° to remain in a nonrotated state. The motion distance from the unannealed 30*S* subunit to the nonrotated and fully rotated states was 10.5 and 8.2 Å, respectively (Fig. 4C). The heated 30*S* subunit exhibited a similar orientation to the unannealed 30*S* subunit and remained in the intermediate state between the nonrotated and fully rotated states (Fig. 4A, C). Distinct from the unannealed and heated 30*S* subunits, the annealed 30*S* subunit adopted an orientation very close to the nonrotated state, with a rotation angle of 2.2° and a maximum motion distance of 5.4 Å (Fig. 4D). The slight difference may be attributable to gentle swirling of the 30*S* head domain. Apparently, annealing switched the 30*S* subunit from an intermediate state to a nonrotated state (Fig. 4E), which is an appropriate state for a new ribosome cycle (34).

The 30*S* subunits under most annealing conditions adopted very similar orientations as the standard annealing condition in the nonrotated states (fig. S10A-F). Even when we extended the hold time after annealing to 24 h at 0°C, we resolved the 30*S* subunit at an average resolution of 3.9 Å and the subunit remained in the nonrotated state, which indicated that the nonrotated state of the 30*S* subunit may be an energetically favorable state. The exception is that when we slowly cooled the heated 70*S* ribosome from a value of 37°C to a value of 4°C over 2 h in the PCR machine, the orientation of the 30*S* subunit was similar to the unannealed and heated ribosomes in the intermediate state (fig. S10G). Considering that the local resolution of the 30*S* subunit from a slowly cooled ribosome was substantially inferior compared with the annealed ribosome, the new orientation imparted by annealing was relevant to the resolution improvement.

### Annealing minimizes the free energy

We used free-energy landscape analysis to investigate the correspondence between the annealing-induced resolution improvement and conformational change. As the major conformational change during polypeptide synthesis, inter-subunit ratcheting between the 50*S* and 30*S* subunits was our proxy to monitor the difference between the unannealed and annealed 70*S* ribosomes. To align the intermediate states, we merged 100,000 particles from the unannealed and 100,000 particles from the annealed ribosomes with different labels into one dataset, and we subjected the merged dataset to a manifold-based analysis in the same conformational coordinate as described elsewhere (35). In the projection direction approximately orthogonal to the interface between the 50*S* and 30*S* subunits, the manifold indicated two clusters, wherein particles from the annealed and unannealed 70*S* ribosomes were well-separated from each other (Fig. 5A, movie S2). This result corresponds to the obvious orientation difference between the unannealed and annealed 70*S* ribosomes.

**Fig. 5.**
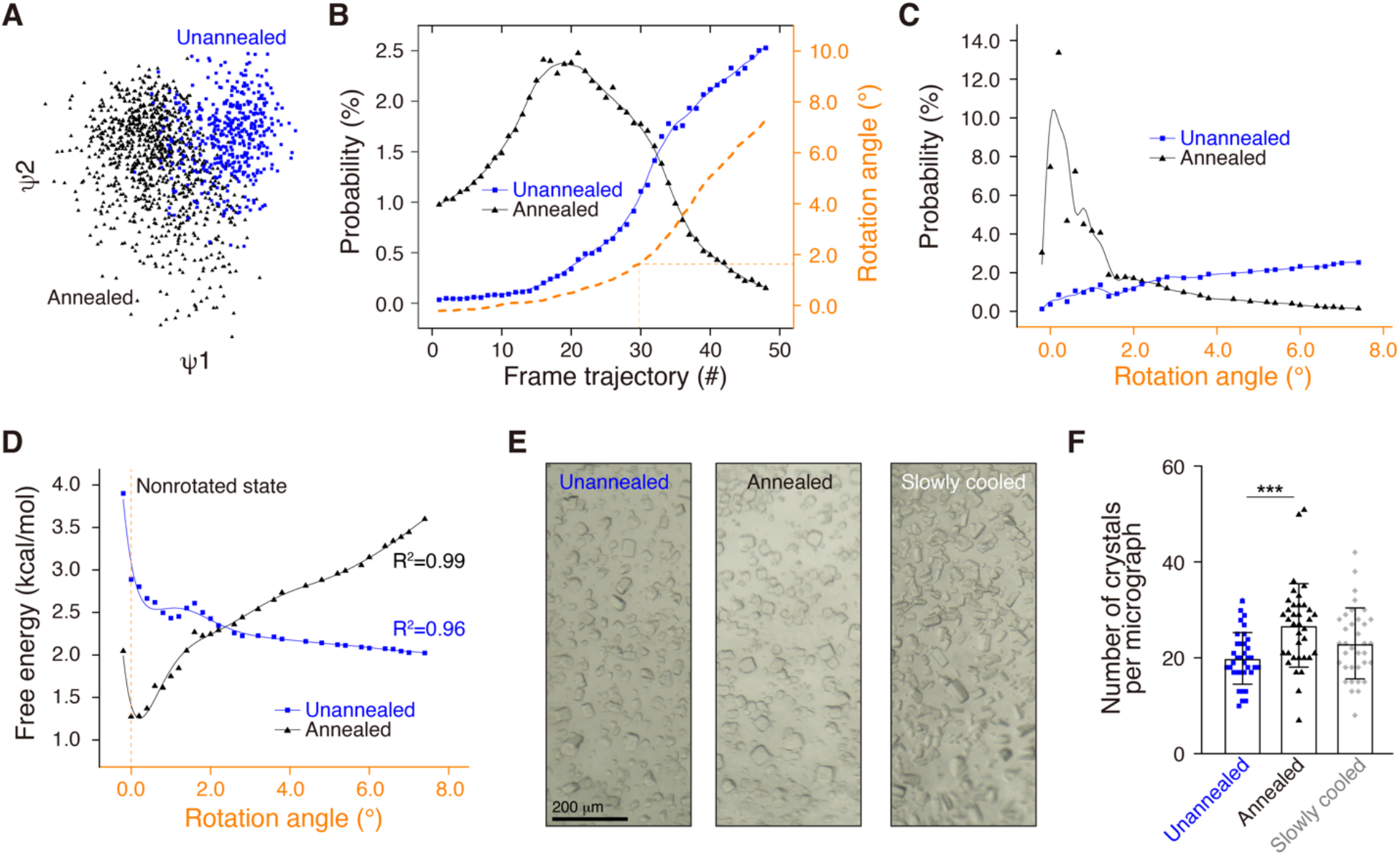
Annealing minimizes the free energy of the 70*S* ribosome. (**A**) Initial manifold snapshots of the 70*S* ribosome in one projection direction (points colored in accordance with the unannealed and annealed subsets). The projection direction is approximately orthogonal to the interface between the 50*S* and 30*S* subunits. (**B**) Particle distribution of unannealed and annealed ribosomes along the frame trajectory. We reconstructed the 3D structure at each frame and calculated the rotation angle of the 30*S* subunit with respect to the nonrotated state. (**C**) Particle distribution of the unannealed and annealed ribosomes along the rotation angle. We recalculated particle numbers in accordance with the rotation angle at intervals of 0.2° and used a moving average to smooth data variation. (**D**) Free-energy distribution of unannealed and annealed ribosomes along the rotation angle. We calculated the energy from the fitted curve in (**C**). (**E**) and (**F**) Growth rate comparison of ribosome crystals: unannealed, annealed, and slowly cooled. We show three representative images for ribosome crystals under three conditions (**E**), and a statistical analysis on the numbers of crystals per micrograph (**F**). We randomly selected 36 images per condition and performed analysis of variance for comparison.

We classified the merged particles into 50 frames based on the orientation of the projections, and we separately calculated the particle number of each frame via the labels corresponding to either the unannealed or annealed ribosomes. The unannealed and annealed 70*S* ribosomes exhibit different peak positions along the frame trajectory. The annealed ribosome was mainly featured in the first 25 frames, whereas the unannealed ribosome started from frame 20 (Fig. 5B). Apparently, the unannealed and annealed ribosomes have distinct particle distributions along the frame trajectory. We reconstructed particles of each frame into a 3D structure with a resolution ranging from ∼5.0–10.0 Å, and we fit the respective atomic models for each frame via rigid body docking of the 50*S* and 30*S* subunits. As the entire body, we aligned the 70*S* ribosome from different frames based on the 50*S* subunit, and we calculated the rotation angles of the 30*S* subunit in different frames relative to the nonrotated state. The rotation angles from frames 1–30 were limited within an ultra-small range of 1.5°, a range that included more than 55% of the particles (the ratio to the total 100,000 particles) of the annealed ribosome. From frames 31–50, the rotation angles of 30*S* subunit were ∼5.0°, a range that included ∼12% of the annealed particles (Fig. 5C). As the control, the unannealed ribosome included 9% of the particles in frames 1–30 and 36% in frames 31–50.

We recalculated the particle distribution of the unannealed and annealed ribosomes against the rotation angles at a step size of 0.2°, and we conducted a moving average to smooth data variations. In the new particle distribution curve, the annealed ribosomes exhibited a sharp peak, which was closer to the nonrotational state; a rotation angle of 0° (Fig. 5C). The sharp distribution indicated improved local resolution with stronger structural rigidity of the annealed ribosome. As the control, the unannealed ribosome exhibited no obvious peaks across the rotation angles from 0° to 8.0°. The wider distribution inevitably deteriorated the local resolution of reconstruction, which fits well with the lower resolution in the unannealed 30*S* subunit.

Based on the Boltzmann distribution, we calculated the free energy of various conformations, including both unannealed and annealed ribosomes, against the rotation angles of the 30*S* subunit. Accordingly, the annealed ribosome exhibited a narrower and deeper well with a minimum energy, whereas the unannealed ribosome exhibited a wider and swallower well. Based on our experimental data, annealing led the 70*S* ribosome from a wider and swallower well (local minima) to a narrower and deeper well (minimum energy state, global minima), and dramatically improved the resolution (Fig. 5D).

Deduced from the Boltzmann distribution formular, one can integrate all atomic arrangements within one conformational state as follows:

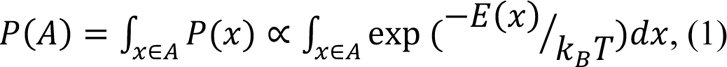

where *P*(*A*) is the probability to remain in conformation *A*. Proteins at lower temperatures have a higher probability of remaining in a narrower and deeper well with a minimum-energy state compared with higher temperatures. Our data quantitatively agrees between predictive theory and experiments and provides direct evidence that annealing synchronizes the 70*S* ribosome at a single-molecule level.

Annealing synchronized the 70*S* ribosome into the homogeneous nonrotated state with minimum energy. We speculate that such ribosomes readily assemble into crystals, as in metallurgy. Actually, crystallization of ribosomes/nucleosomes features an annealing-like treatment, wherein researchers usually warm ribosomes/nucleosomes to between 37°C and 55°C, then decrease to room temperature (19°C) (36, 37). To test assembly into crystals, we performed strict annealing on the 70*S* ribosome and cooled the annealed ribosome to 4°C for crystallization (Fig. 5E). The unannealed ribosome and the slowly cooled ribosome were controls. Statistical analysis showed that the annealed ribosomes formed more crystals than the unannealed and slowly cooled ribosomes. Annealed ribosomes remained in a more homogeneous minimum-energy state, which facilitated assembly into crystals (Fig. 5F).

As the intermediate state of the annealing, we investigated the structural switch from a heated ribosome to an annealed ribosome via free-energy landscape analysis on the combined heated and annealed ribosomes. Similar to the combination of unannealed and annealed ribosomes, heated and annealed ribosomes also exhibited similar particle distribution patterns (fig. S11A-D). As the control, when we analyzed the unannealed and heated ribosomes, the particle distribution was similar, and the heated ribosome had an even wider distribution (fig. S11E). These results indicate that the conformational change occurred during cooling.

## DISCUSSION

Researchers widely use annealing on metals, alloys, and semiconductors to decrease the number of defects in crystals and increase the size of the crystals. Here, we experimentally demonstrated that annealing can also synchronize the 70*S* ribosome into the minimum-energy state on a single-molecule level (Fig. 6). After annealing, the 70*S* ribosome readily assembled into 3D crystals, which will inspire researchers to further investigate the physical and chemical effects of annealing on the 70*S* ribosome and other biological macromolecules.

**Fig. 6.**
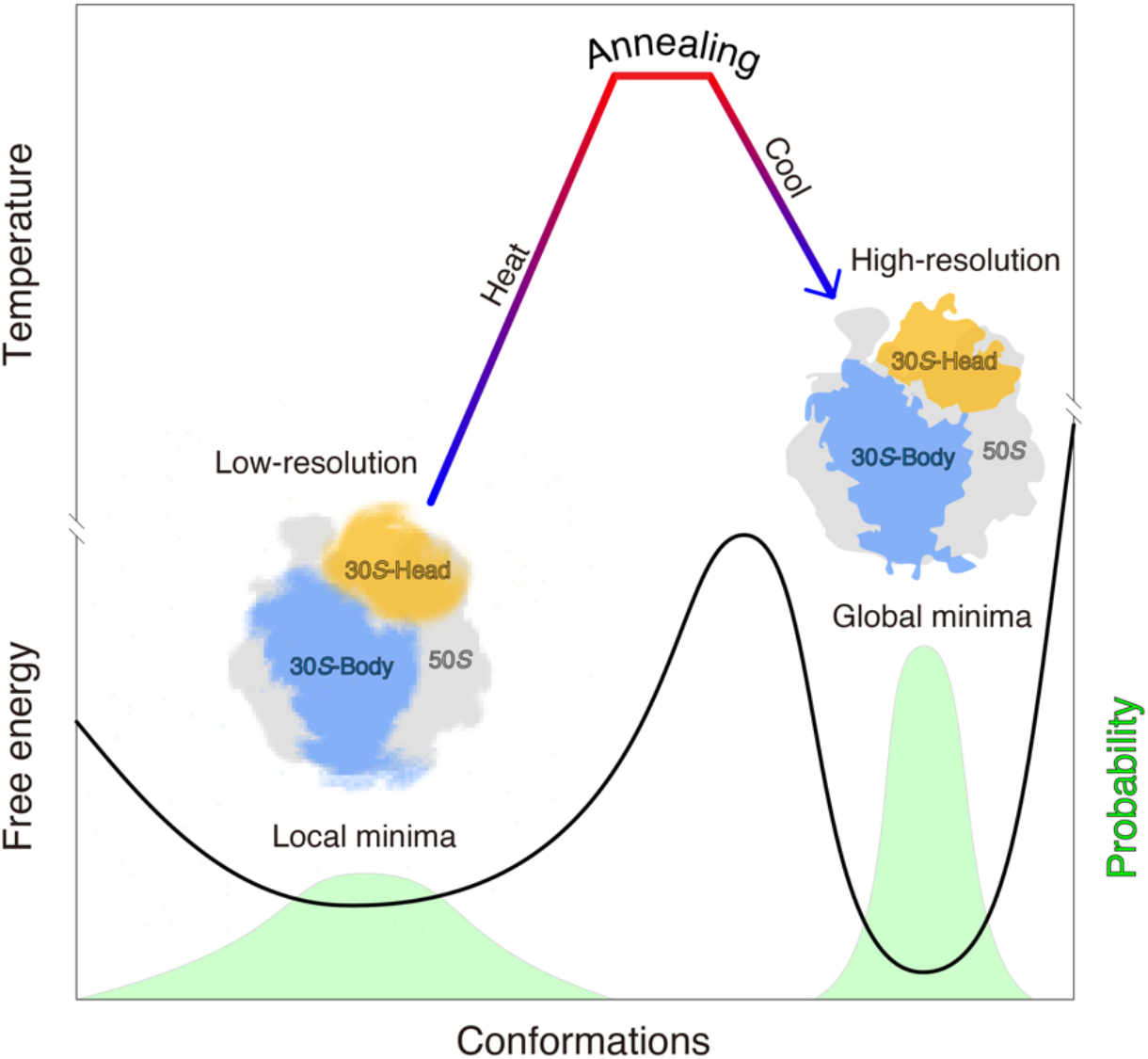
Model illustrating that annealing can synchronize a ribosome into a minimum-energy state with improved local resolution. Both the free-energy curve (solid line) and particle distribution probability (light green peaks) are shown.

Structural flexibility, albeit critical for protein function, hinders researchers’ ability to apply structural studies to elucidate function on a molecular level. Sustained efforts—such as mutations on key residues (38, 39), introduction of additional disulfide bonds (40, 41), addition of antibody/binding proteins (42–44), or cross-linking in a solution or within a glycerol/glucose gradient (45, 46)—have been useful for optimizing samples to increase structural stability. However, such efforts are time-consuming and can lack clear direction, and the final structures are confined to fixed states and sometimes are even distorted after additional manipulations. Annealing—a combination of appropriate heating and cooling—is non-destructive to proteins, and is a facile yet robust approach for high-resolution cryo-EM.

Researchers have also attempted to improve resolution via local refinement on flexible regions during cryo-EM image processing (47–50). We performed local refinements on the flexible 30*S* subunit on both unannealed and annealed ribosomes. The average local resolution of the 30*S* subunit of the unannealed 30*S* subunit was improved ∼1 Å to a value of 4.2 Å after local refinements (figs. S12 and S13). Distinct from the resolution improvement via annealing, local refinement in itself still resulted in an insufficient average resolution of 5.5 Å on the head domain of the 30*S* subunit (figs. S12 and S13). Apparently, annealing and local refinement improve local resolution via different mechanisms. Annealing can drive biological macromolecules into a minimum-energy state and globally increase the resolution across the entire map, irrespective of the region size. As a control, local refinement worked on an algorithmic level and was only applicable to regions with reasonable sizes. When we applied local refinement to the annealed ribosome, both the body and head domains of the 30*S* subunit were improved to 2.9 and 3.9 Å, respectively (figs. S12 and S13). This indicates that annealing is compatible with local refinement on flexible regions and can further optimize local resolutions for detailed structural analysis.

One can use annealing to synchronize proteins into a minimum-energy state. Analogous to cell synchronization, protein/conformation synchronization on a single-molecule level may be helpful for many single-molecule approaches, such as those which use optical tweezers and single-molecule fluorescence resonance energy transfer (51–53). One can also use annealing to investigate temperature adaptation and protein folding, and to facilitate algorithm development in molecular dynamics simulation. Thus, researchers should thoroughly investigate the mechanism of annealing and further optimize the annealing conditions for improved resolution.

## Acknowledgments

We thank Prof. Yifan Cheng from UCSF for thorough discussions. We thank Prof. Abbas Ourmazd and Dr. Ghoncheh Mashayekhi from UW-Milwaukee for help on ManifoldEM. We are grateful to Kang Li, Dianli Zhao and Ceng Gao from the CryoEM facility for Marine Biology at QNLM for our cryo-EM data collection.

## Funding

National Key R&D program of China, 2017YFA0504800 (QS)

National Key R&D program of China, 2018YFC1406700 (QS)

National Natural Science Foundation of China, 31870743 (QS)

## Author contributions

Conceptualization: QS

Methodology: XC, XS, LQ, GL, YL, JL

Investigation: XC, XS, ML, LL, TL, YQ, GL

Visualization: XC, XS, QS

Funding acquisition: QS

Project administration: QS

Supervision: QS

Writing – original draft: XC, XS, LL, QS

Writing – review & editing: QS

## Data availability

The cryo-EM density maps of ribosomes under all conditions (S1-S10) were deposited in Electron Microscopy Data Bank (EMDB) with the accession numbers from 31266 to 31275, as listed in fig. S2B. The locally refined cryo-EM maps of the 30*S* subunits from the unannealed (S1) and annealed (S5) ribosomes were also deposited in EMDB with the respective accession numbers of 31277 and 31279. Cryo-EM density maps and models for all 50 states in free energy landscape analysis are available from the corresponding author by request. All other data is available in the main text or the supplementary materials.

## Code availability

The codes for local resolution analysis and modified ManifoldEM have been uploaded to GitHub: https://github.com/soothing35/cryoEM_annealling.

## Supplementary Materials

### Materials and Methods

#### Purification and assembly of the apo-state 70*S* ribosome

70*S* ribosomes were extracted from *Escherichia coli* KC6/ΔsmpB/ΔssrA/S1::S1-8xHis stain as previously described (1, 2). In brief, cells were lysed and the debris was removed via centrifugation. The supernatant was layered on the top of a sucrose cushion and centrifuged at 29,400 rpm using a 45Ti rotor (Beckman, USA) for 16 h at 4°C for the crude 70*S* ribosomes. The samples were further applied to a Butyl-650S column and eluted with a linear 1.5–0 M (NH_4_)_2_SO_4_ gradient in Tris-HCl (20 mM, pH 7.4) buffer containing 400 mM KCl, 10 mM Mg(OAc)_2_, and 6 mM 2-Mercaptoethanol. The pooled fractions containing pure ribosomes were further passed through a 5 mL HisTrap Excel column (GE Healthcare Life Sciences, USA) to remove residual ribosomes with ribosomal protein S1 bound. The S1-free 70*S* ribosome was loaded on 10%–40% (w/v) sucrose gradient using SW32 rotor (Beckman, USA) by centrifugation at 23,000 rpm for 14 h at 4°C. The fractions containing 70*S* ribosomes were pooled and dialyzed against Tris-HCl (5 mM, pH 7.4) buffer containing 60 mM NH_4_Cl, 10 mM MgCl_2_, and 6 mM 2-Mercaptoethanol. The dialyzed ribosomes were flash-frozen in liquid nitrogen, and stored under −80°C.

The same batch of 70*S* ribosomes was used for cryo-EM sample preparation and image processing under various conditions.

#### Cryo-EM sample preparation

The frozen 70*S* ribosomes were melted on ice and then diluted to a final concentration of 700 nM in HEPES (5 mM, pH 7.5) buffer containing 50 mM KCl, 10 mM NH_4_Cl, 10 mM Mg(OAc)_2_, and 2 mM 2-Mercaptoethanol. To ensure fast heat exchange, no more than 20 µL of the diluted ribosomes were pipetted into one PCR tube. All the samples were subjected to various treatments as listed in Fig. 2B, before vitrification.

For the annealed treatments (S4–S10), the diluted ribosomes were placed on ice for 5 min and then transferred to a water bath, with preset temperatures ranging from 30°C to 65°C. After a 1- or 5-min hold time, the heated samples were immediately immersed into a mixture of salt, ice, and water (measured temperature: −18°C) for <5 s; and then to a mixture of ice and water (measured temperature: −0°C), or directly immersed into a mixture of ice and water for either 5 min or 24 h. As a control, the unannealed ribosome (S1) was simply immersed in a mixture of ice and water for 5 min. For the slowly annealed sample (S3), the diluted ribosome in the PCR tube was placed in a PCR machine (Bio-Rad, USA). The temperature decreased gradually from an initial value of 37°C to a final value of 4°C over 2 h at a cooling rate of 0.275°C/min.

After treatments, all the unannealed and annealed samples in the PCR tubes were immediately placed in a Vitrobot chamber set at the minimum temperature of 4°C and a humidity of 100% (Mark IV, Thermo Fisher Scientific, USA). A gold-supported holey carbon grid was glow discharged and pre-cooled in the Vitrobot chamber for 2 min, and 4 µL of solution was applied onto the grid. The grid was immediately plunged into liquid ethane and stored under liquid nitrogen for future cryo-EM imaging.

#### Cryo-EM sample preparation with a mist umbrella

In temperature-resolved cryo-EM, sudden exposure of high-temperature mist (∼37°C) from a Vitrobot chamber to cooler ambient temperatures (∼20°C) causes substantial ice contamination. To minimize ice contamination under various temperatures, especially at high temperatures, a mist umbrella was designed. The mist umbrella was made of a piece of round filter paper (40-mm diameter) with a precut square hole (9.0 mm × 3.6 mm) prepared via a home-made device (fig. S7). The peripheral belt of the mist umbrella was pre-treated with low-adhesive glue. Before lifting the Vitrobot tweezer into the chamber, the mist umbrella was clamped to the tweezer via the precut hole. Over the course of lifting the grid to the chamber, the mist umbrella fell off the tweezer yet glued to the bottom surface of the chamber. After blotting, the shutter of the Vitrobot chamber was immediately opened and the grid passed through the precut hole into liquid ethane. The high-temperature mist in the chamber was blocked by the mist umbrella, which prevents the exposure of high-temperature mist to the cool ambient. When the liquid nitrogen reservoir and cryogenic cup dropped away from the chamber, the tweezer reattached the mist umbrella to shield the grids (movie S1). During grid transfer from liquid ethane to liquid nitrogen, one can readily remove the mist umbrella from the Vitrobot tweezer along the tear line.

The mist umbrella was used for the heated ribosome (S2) as an intermediate state. Diluted ribosomes were incubated in a water bath at 37°C for 5 min, and then transferred into the Vitrobot chamber at a temperature of 37°C and a humidity of 100%. The grid was placed in the chamber for at least 100 s to ensure complete heat exchange, and then 5 µL of solution was applied onto the grid via pre-heated tips. The grids were blotted with a 1 s blotting time and force level of 2, immediately plunged into liquid ethane, and stored under liquid nitrogen for future cryo-EM imaging.

#### Cryo-EM data collection

Cryo-EM grids were examined in a low-dose mode on a Talos L120C TEM (Thermo Fisher Scientific, USA) for screening. Data collection on high-quality grids in all conditions was performed with the same Titan Krios G^3i^ microscope (Thermo Fisher Scientific, USA) equipped with a K3 BioQuantum direct electron detector (Gatan, USA). Special care was taken to perform a coma-free alignment on the microscope.

Movies were collected via FEI EPU (3) automated data collection software at a total dose of 60 e^−^/Å^2^ fractionated over 50 frames in a defocus range of −1 to −1.5 µm. A super-resolution mode was used with a final pixel size at 0.53 Å. The numbers of the collected movies under different conditions are listed in fig. S2B and table S1.

#### Cryo-EM data processing and 3D reconstruction

For optimal structural comparison, the same image processing strategy was performed on ribosomes under all conditions with cryoSPARC v.3.1.0 (4); fig. S2 shows a detailed workflow. Specifically, raw movie stacks were aligned and summed in accordance with dose weighting with MotionCor2.1 (5). CTF parameters of the summed micrographs were determined with CTFFIND-4 (6). Micrographs with maximum resolution estimates of at least 5 Å were imported into cryoSPARC. Automatic particle picking was performed on the selected micrographs, and particle sets were created and subjected to reference-free 2D classifications. Obvious junk was excluded from the particle set. Six reference structures—including 2 × 70*S* ribosome (PDB ID 5MDZ), 2 × 50*S* subunit (derived from PDB ID 5MDZ), and 2 × 30*S* subunit (derived from PDB ID 5MDZ)—were low-pass filtered to 40 Å and utilized for heterogeneous refinement. Only particles in the integral 70*S* ribosome with sufficient signal on the 30*S* subunit were collected. Two hundred thousand particles were randomly selected from the new particle set for each condition, and per-particle refinement of CTF parameters was conducted. Particles with local CTF parameters were subjected to homogeneous refinement. All the final reconstructions were filtered and sharpened by a cryoSPARC post-processing session and the resolutions were determined by gold-standard FSC 0.143.

#### Model building and structural analysis

The crystal structure of the 70*S* ribosome, with an empty A site (PDB ID 5MDZ) (7), from *E*. *coli* was chosen as the initial template; and mRNA and fMet–tRNA were removed before fitting into the apo-state 70*S* ribosomes. The 50*S* and 30*S* subunits of the modified crystal structure were separately fit into the 70*S* ribosomes under various conditions via rigid-body docking, and the fits were ideal except on the L1 stalk. A flexible fitting of the L1 stalk was performed with Rosetta 2018.33.60351 (8). The atomic model was further optimized for improved local density fitting with Coot 0.8.9.1 (9) and real-space refinement with PHENIX 1.17.1 (10).

Structures of the unannealed, heated, and annealed 70*S* ribosomes were aligned using their 50*S* subunits as a reference. The conformational change of the 30*S* subunits was analyzed in Pymol 2.4.0 as described elsewhere (11, 12). Difference vectors between the phosphorous and C*α* atoms in the unannealed, heated, and annealed 30*S* subunits were calculated via the modified script (https://github.com/soothing35/cryoEM_annealling/Pymol_scripts), and rotation angles between the 30*S* subunits were measured with the Pymol script ‘RotationAxis.py’ (https://pymolwiki.org/index.php/RotationAxis).

Other structural analysis, such as structural superimposition, were fulfilled with UCSF Chimera 1.16 (13).

#### Local refinement on the 30*S* subunit

Local refinement on the 30*S* subunit was also performed with cryoSPARC (4). Based on the docked atomic model, a mask for the 50*S* subunit was created and applied to subtract the corresponding densities in all of the particles. Local refinement was conducted on the 30*S* subunit density with rotations and shifts re-centered every iteration. A new function in cryoSPARC termed “using pose/shift gaussian prior during alignment” was also used for improved reconstruction. The final reconstruction was filtered and sharpened by a cryoSPARC post-processing session and the resolutions were determined by gold-standard FSC 0.143.

#### Local resolution quantification

To compare local resolutions of the 30*S* subunits and other sub-regions under various conditions, local resolution quantification was calculated via averaging values of the local resolution in selected regions (fig. S4). Specifically, local resolutions on the entire 70*S* ribosomes under various conditions were first estimated as per local resolution estimation with cryoSPARC (14). Then the regions within a certain local resolution range or within the same functional domain were selected as the targets. After atomic model building, a mask corresponding to the targeted region was created with the extension of 2 pixels, and the same mask strategy was applied to the targeted regions from other structures. The resulting sub-regions were adjusted to the same volume among different structures, and the values of the local resolution within the mask were read via our Python script (https://github.com/soothing35/cryoEM_annealling/locres_points). The numbers of local resolution values in the range of 2.0–2.1 Å, 2.1–2.2 Å, and so on were counted in different sub-regions. The respective means and standard deviations were calculated for each sub-region and compared with each other.

#### Free-energy landscape analysis

Free energy is inversely proportional to particle distribution, as described by the Boltzmann distribution (15). The particle distribution of 70*S* ribosomes under various conditions was calculated with ManifoldEM (https://github.com/GMashayekhi/ManifoldEM_Matlab) as described elsewhere (16, 17). Specifically, 100,000 particles were randomly selected from screened particle sets for each condition to sample the entire conformational trajectory. Particles files, including both .star and .mrcs, were selected after auto-refinement with RELION 3.0.8 (18). To ensure accurate alignment of structures from two different conditions, their particle files were merged together via appending one to the other with different labels, and the merged particle files were analyzed with ManifoldEM for further analysis.

In ManifoldEM, merged particles were classified in accordance with the pre-determined Euler angels, and particle images were deemed as dots in high-dimensional space. Clouds of dots in high-dimensional space were eigen-decomposed and embedded into low-dimensional space with a diffusion map algorithm, where the kernel was defocus-tolerant. The first five eigen-vectors were selected and the conformational changes along each eigen-function were extracted into 2D movies by nonlinear Laplacian spectral analysis (NLSA), also termed non-linear singular value decomposition (SVD). Considering that the inter-subunit rotation between the 50*S* and 30*S* subunits was considered to be the primary conformational change, movies with such conformational changes along each eigen-function were chosen manually. Particles involved in the conformational changes were further classified into 50 classes along the conformational trajectory.

For all 50 classes, 3D volumes were reconstructed via back-projection in RELION 3.0.8. Atomic models for the 50*S* and 30*S* subunits were separately docked in each volume via rigid-body docking. All structures were aligned to the nonrotated state (PDB ID 4V9D) with the 50*S* subunit as a reference. The rotation angle of the 30*S* subunit in each class was calculated with respect to the 30*S* subunit in the nonrotated state. Based on the labels of different conditions, particles in each class were separated. The particle distribution within the same condition was recalculated and the respective free energy was calculated based on the Boltzmann distribution.

#### Ribosome crystallization

*E. coli* 70*S* ribosomes were readily crystalized when their native ribosomal protein L9 was replaced with *Thermus thermophilus* ribosomal protein L9 (19). *E. coli* L9-free 70*S* ribosomes were expressed in *E. coli* KC6/ΔsmpB/ΔssrA/S1::S1-8xHis/EcoL9(40)FRT strain and purified as described above. *T. th* ribosomal protein L9 was expressed in *E. coli* and obtained through multiple purification steps, including heating at 70°C to denature *E. coli* endogenous proteins, anion exchange chromatography via Mono S HR 16/10, and gel filtration chromatography in HiLoad 26/600 Superdex 75pg. Subsequently, the purified *T. th* L9 protein were concentrated and stored in Tris-HCl (10 mM, pH 7.4) buffer containing 50 mM KCl and 1 mM 2-Mercaptoethanol.

*E. coli* L9-free 70*S* ribosome at a final concentration of 4 µM was incubated with 8 µM *th* L9 on ice bath in the HEPES buffer. The mixtures were split into three portions and each portion was subjected to various annealing treatments. One portion of proteins was fast immersed into the mixture of salt, ice and water after heating to 37°C, and then placed on ice bath (depicted as the annealed state); one portion of proteins was heated to 37°C and then slowly cooled down to 4°C over a period of 2 hours (depicted as the slowly cooled state); the third was kept on ice bath as the control.

The same crystallization conditions were applied to all annealing treated samples. Briefly, 3 µL of sample and equal volume of precipitation (2.9% (w/v) PEG 20000, 100 mM Tris-HCl, pH 7.4, 10% (v/v) MPD, 160 mM L-Arginine-HCl, and 0.5 mM 2-Mercaptoethanol) were mixed and crystallized in a 24-well sitting drop plate containing 500 µL of reservoir at 4°C for 14 days. For each condition, 36 images from seven drops were randomly captured, and crystals larger than 20×20×40 µm^3^ were counted. The statistical analysis was fulfilled in GraphPad Prism 8.0.

### Supplementary Text

#### Structural comparison of 70*S* ribosomes under various conditions

In our work, 70*S* ribosomes under various conditions were resolved via single-particle cryo-EM. Corresponding structural comparisons indicated that annealing could synchronize 70*S* ribosomes, especially flexible regions, into homogenous states with improved local resolution. To objectively compare 70*S* ribosomes, we tested reconstruction as follows:

1) Same batch of ribosomes. 70*S* ribosomes were aliquoted and stored at **−**80°C and unused ribosomes were discarded after thawing.
2) Same data collection conditions. The same microscope (Titan Krios G^3i^) with the same camera (Gatan K3 Bioquantum) was used for all data collections. Total electron dose, defocus, and other pertinent values were set to be the same.
3) Same data processing pipeline. The procedure as described in fig. S2A was consistently applied to all 70*S* ribosomes under various conditions.
4) Same data screening strategy. Only obvious junk and disassembled ribosomes were removed from the dataset. In most datasets, 82.3% to 93.7% of the particles were selected after 2D and 3D classifications. Three exceptions were the heated ribosome (S2) and annealed ribosomes (S9 and S10). Ribosomes (S9 and S10) were heated to 55°C and 65°C, respectively (closer to the ∼72°C melting temperature of bacterial ribosome), for 5 min; these 70*S* ribosomes readily fell apart. Regarding S2, ribosomes were vitrified at 37°C as the heated state. Compared with ribosomes under other conditions, more unsatisfactory particles (∼46.9%) were excluded from the final reconstruction, but the local resolution of 30*S* subunit was still inferior.
5) Same number of particles (200, 000) for the final reconstruction, for comparison.
6) Repeating experiments. Repeats were performed on three representative conditions, including the unannealed (S1), heated (S2), and annealed (S5) ribosomes. A new batch of 70*S* ribosomes was used and reconstructed. There was no obvious structural difference between two parallel experiments.

In summary, the structural differences should be attributable to differences in treatments.

**Fig. S1.**
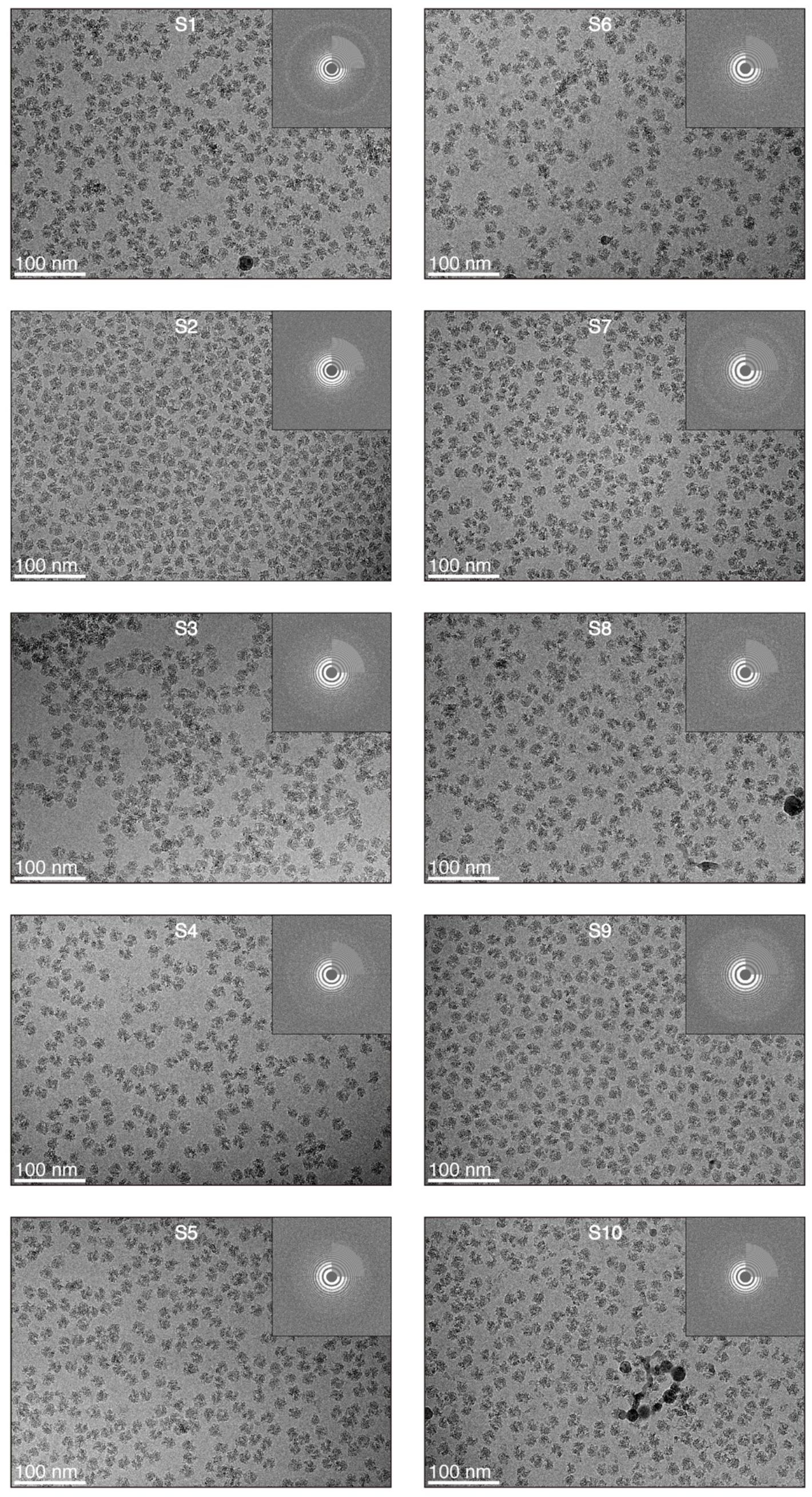
Typical cryo-EM images under various conditions and their respective power spectra. Fig. 2B lists detailed conditions for samples S1–S10.

**Fig. S2.**
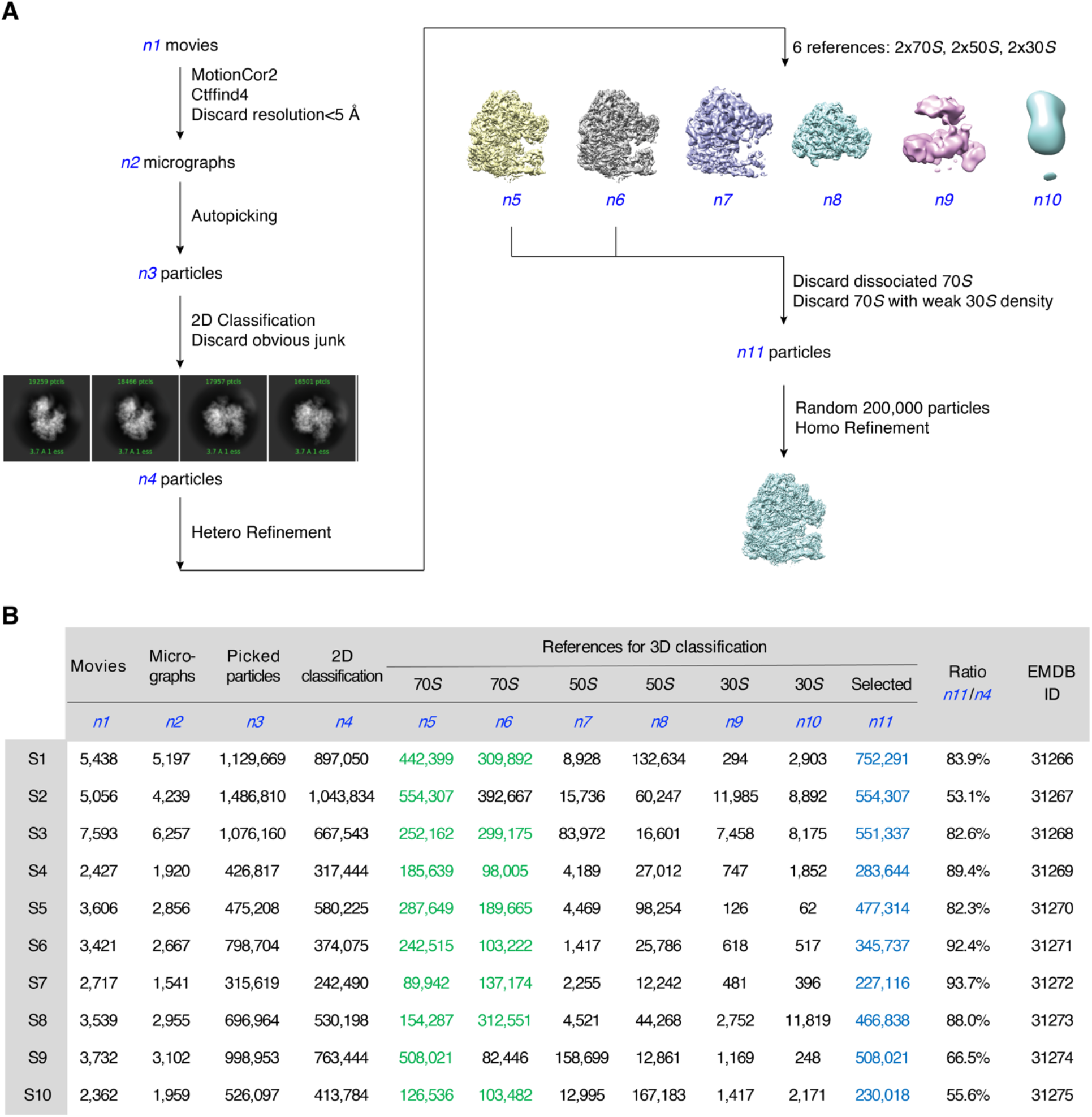
3D reconstructions of 70*S* ribosomes under various conditions. (**A**) The same flow chart was used for 3D reconstruction of 70*S* ribosomes under various conditions. (**B**) Detailed numbers from *n1*–*n11* in (**A**) and the corresponding deposited EMDB IDs are listed.

**Fig. S3.**
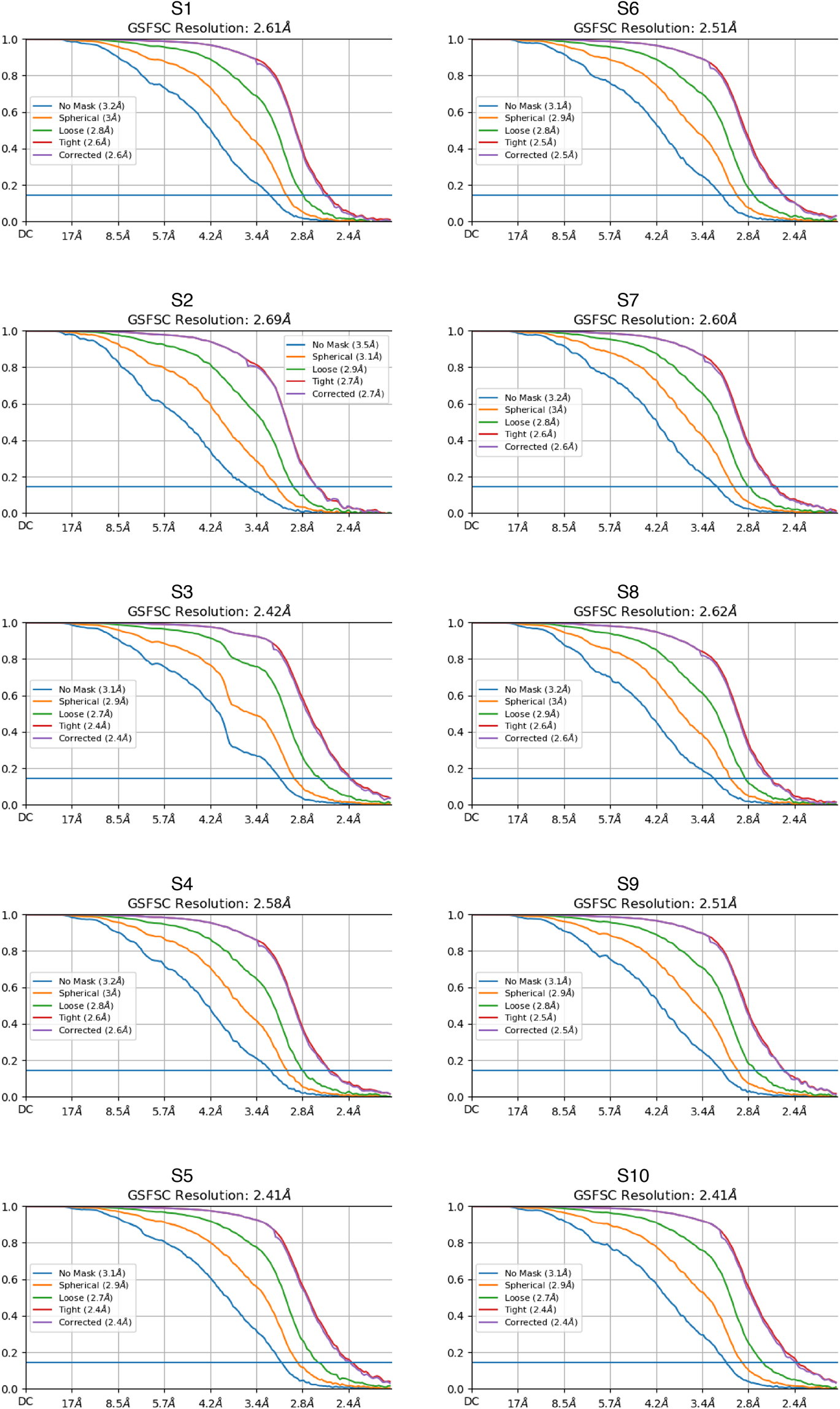
FSC curves for 70*S* ribosomes under various conditions.

**Fig. S4.**
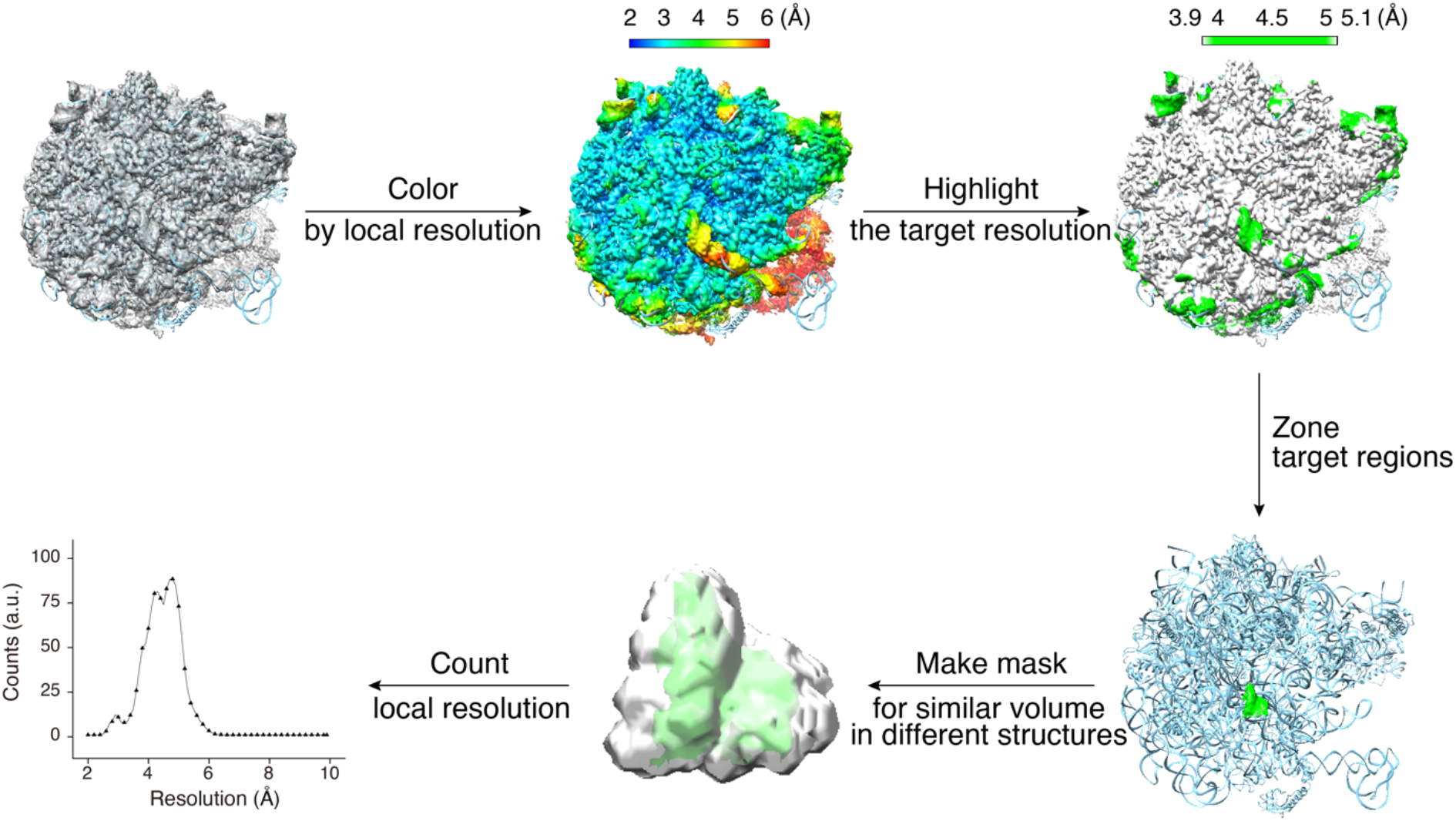
Flow chart for local resolution calculation on selected regions. To compare local resolutions from different structures, the volumes of the target regions in different EM maps were adjusted to the same level via changing thresholds.

**Fig. S5.**
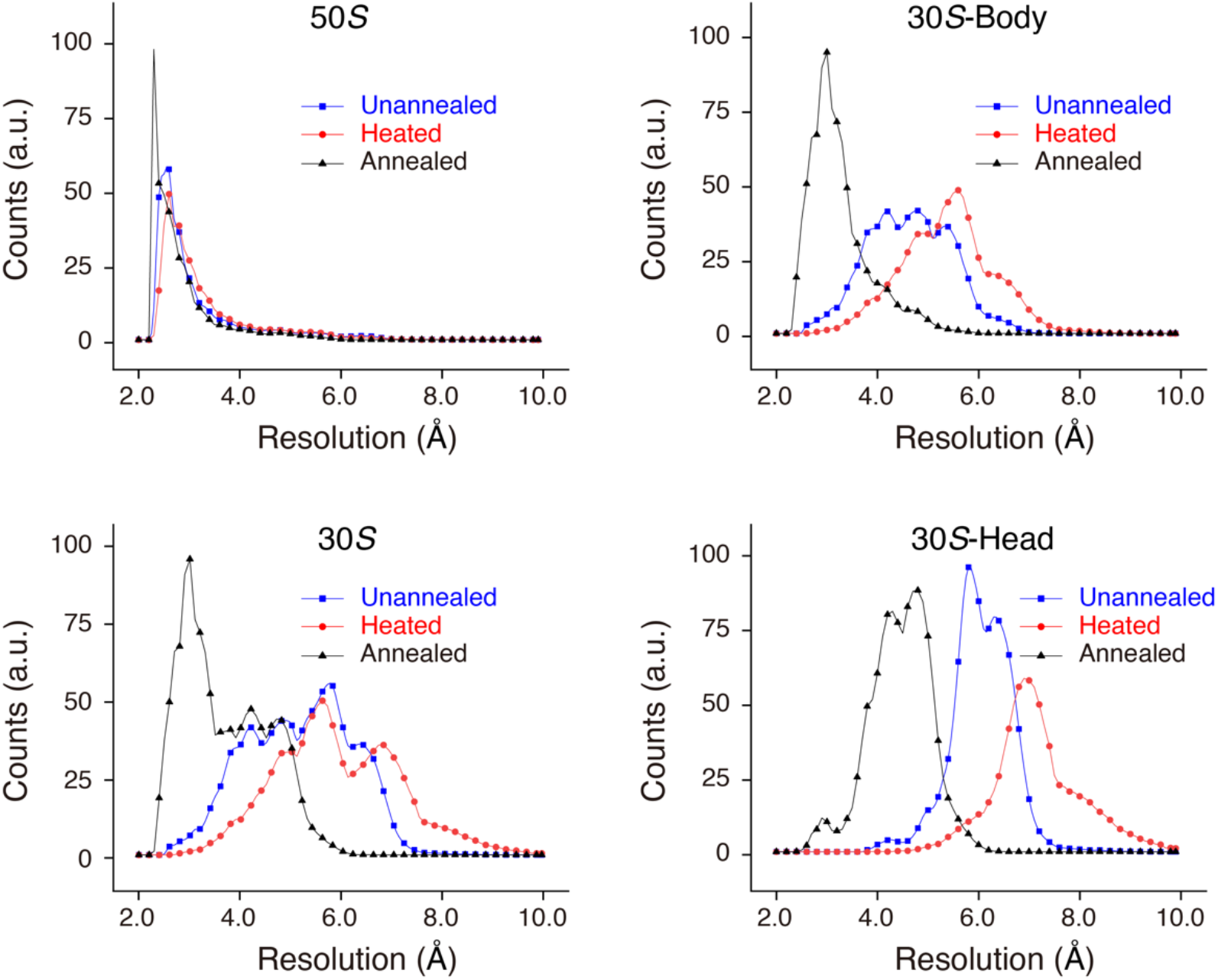
Local resolution histograms of different subdomains from the unannealed, heated, and annealed 70*S* ribosomes. The mean and standard deviation from these histograms were calculated (Fig. 1B). “a.u.” means arbitrary units.

**Fig. S6.**
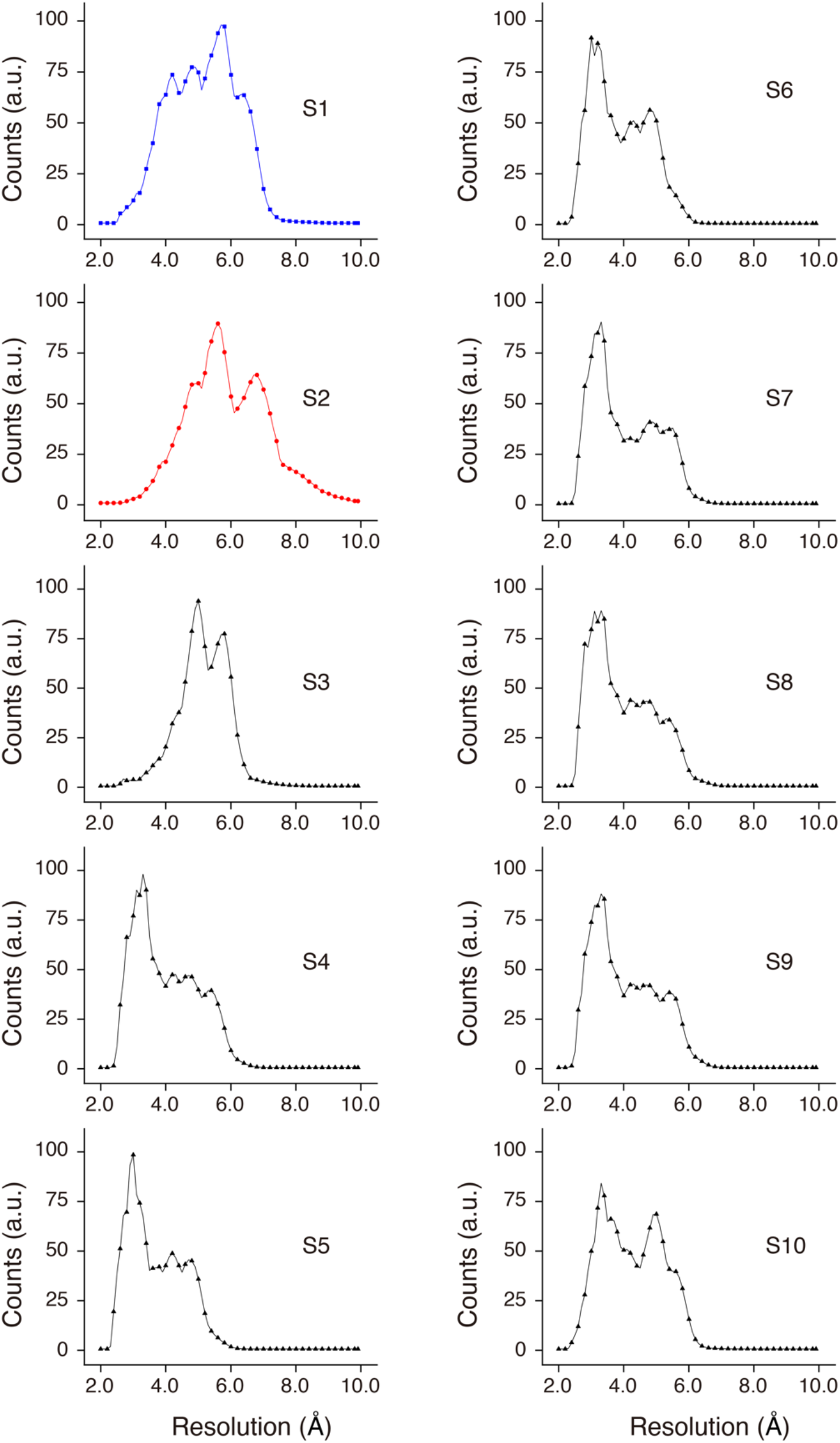
Local resolution histograms of 30*S* subunits under various conditions. The mean and standard deviation from these histograms were calculated (Fig. 2C).

**Fig. S7.**
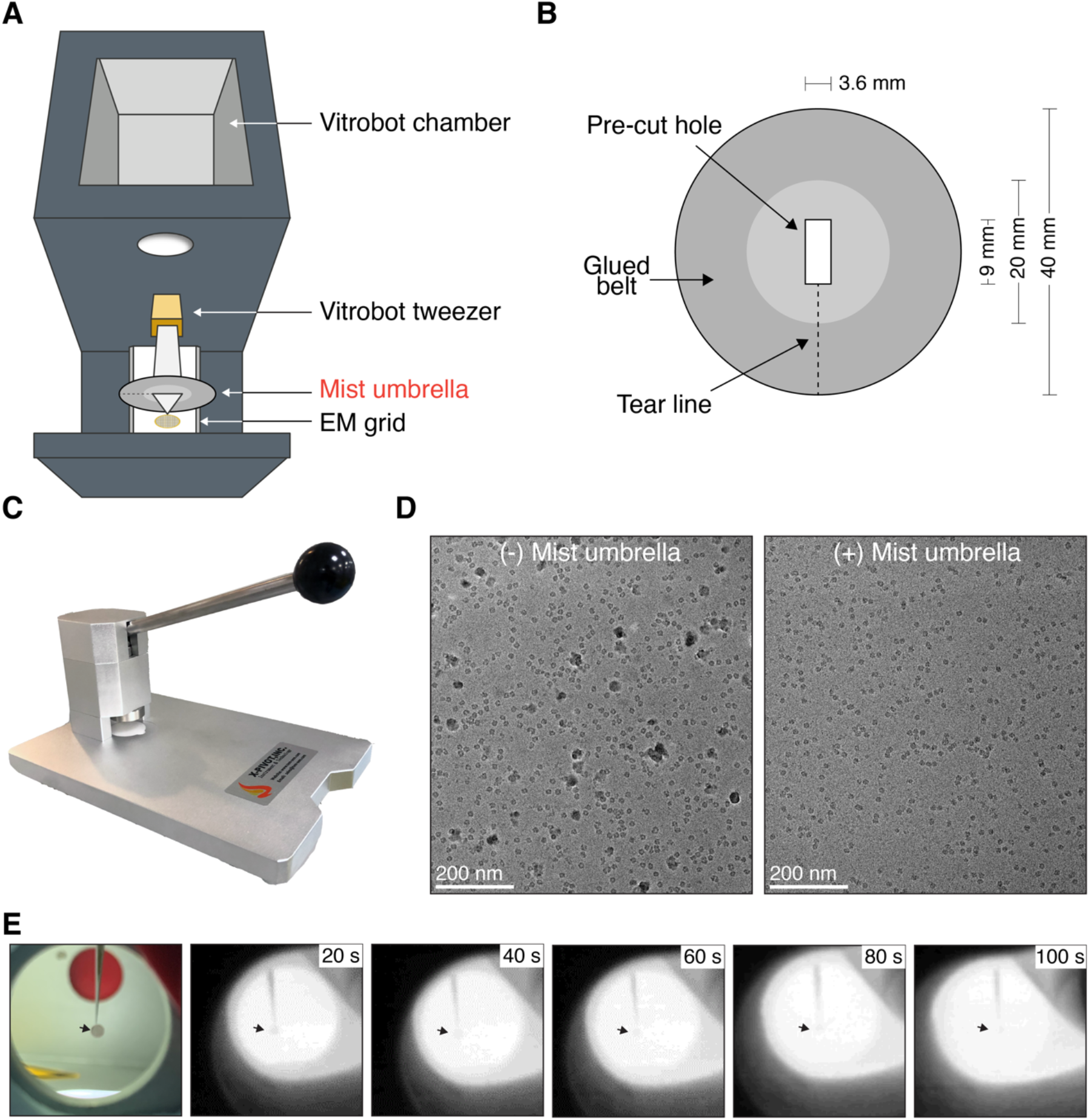
Optimization for temperature-dependent cryo-EM. (**A**) Diagram for the Vitrobot device with a mist umbrella. (**B**) Detailed parameters for the mist umbrella. Briefly, there is a pre-cut hole on round filter paper. (**C**) Home-made device to fabricate mist umbrellas. (**D**) Cryo-EM micrographs with or without a mist umbrella. (**E**) Heat exchange between grids and the Vitrobot chamber reached a balance in ∼100 s. Black arrow points to the position of the cryo-EM grid.

**Fig. S8.**
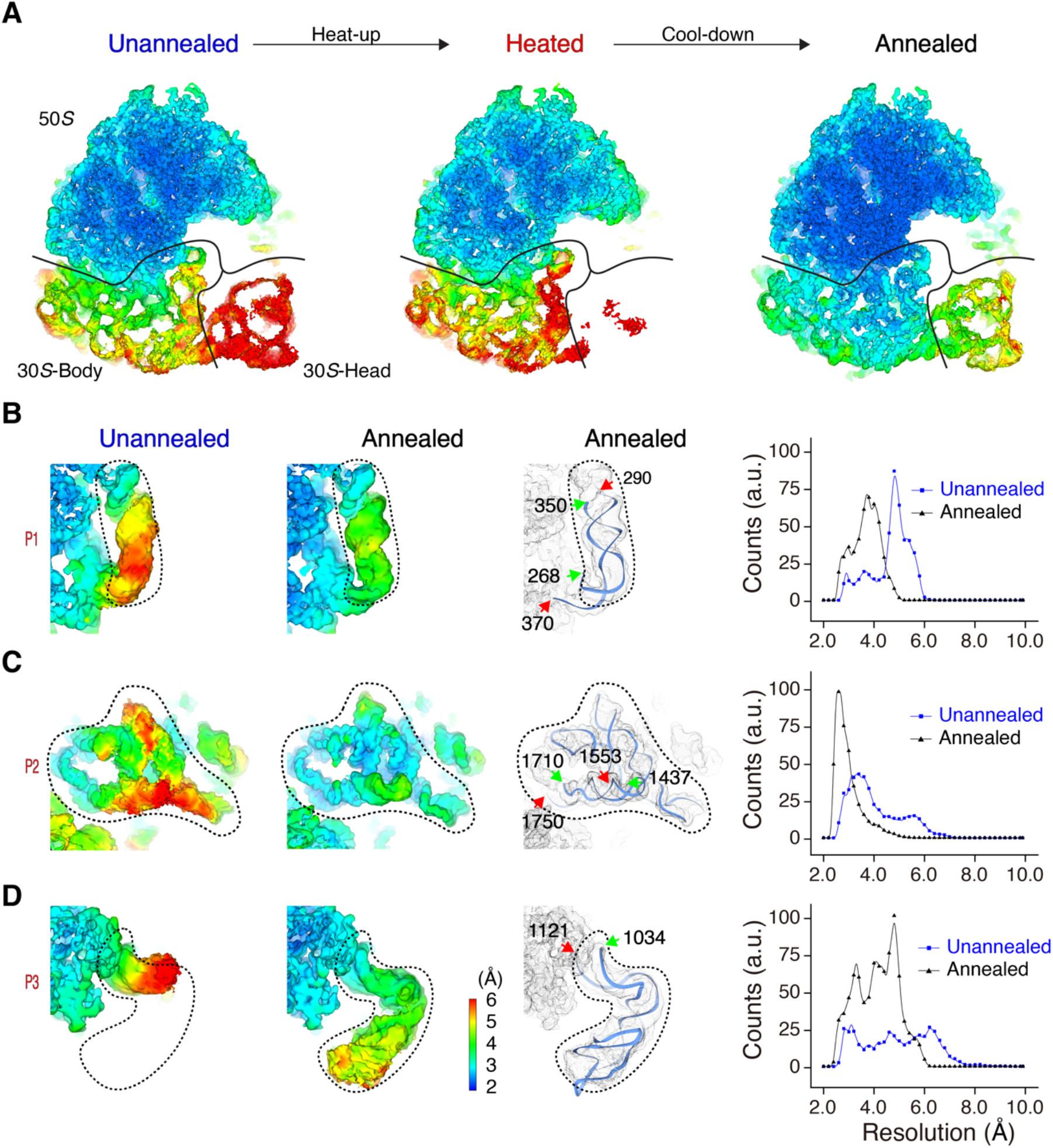
Annealing stabilizes flexible regions on the periphery of the 50*S* subunit. (**A**) Central slice of local resolution maps of the unannealed, heated, and annealed 70*S* ribosomes. (**B**–**D**) Annealing stabilizes three typical flexible regions on the periphery of the 50*S* subunit. The positions of P1, P2, and P3 in 70*S* ribosomes are shown in Fig. 1A. Local resolution maps and histograms in the unannealed and annealed conditions are shown. Models were based on the annealed structures. Green and red arrows indicate the start and end, respectively, of selected strands.

**Fig. S9.**
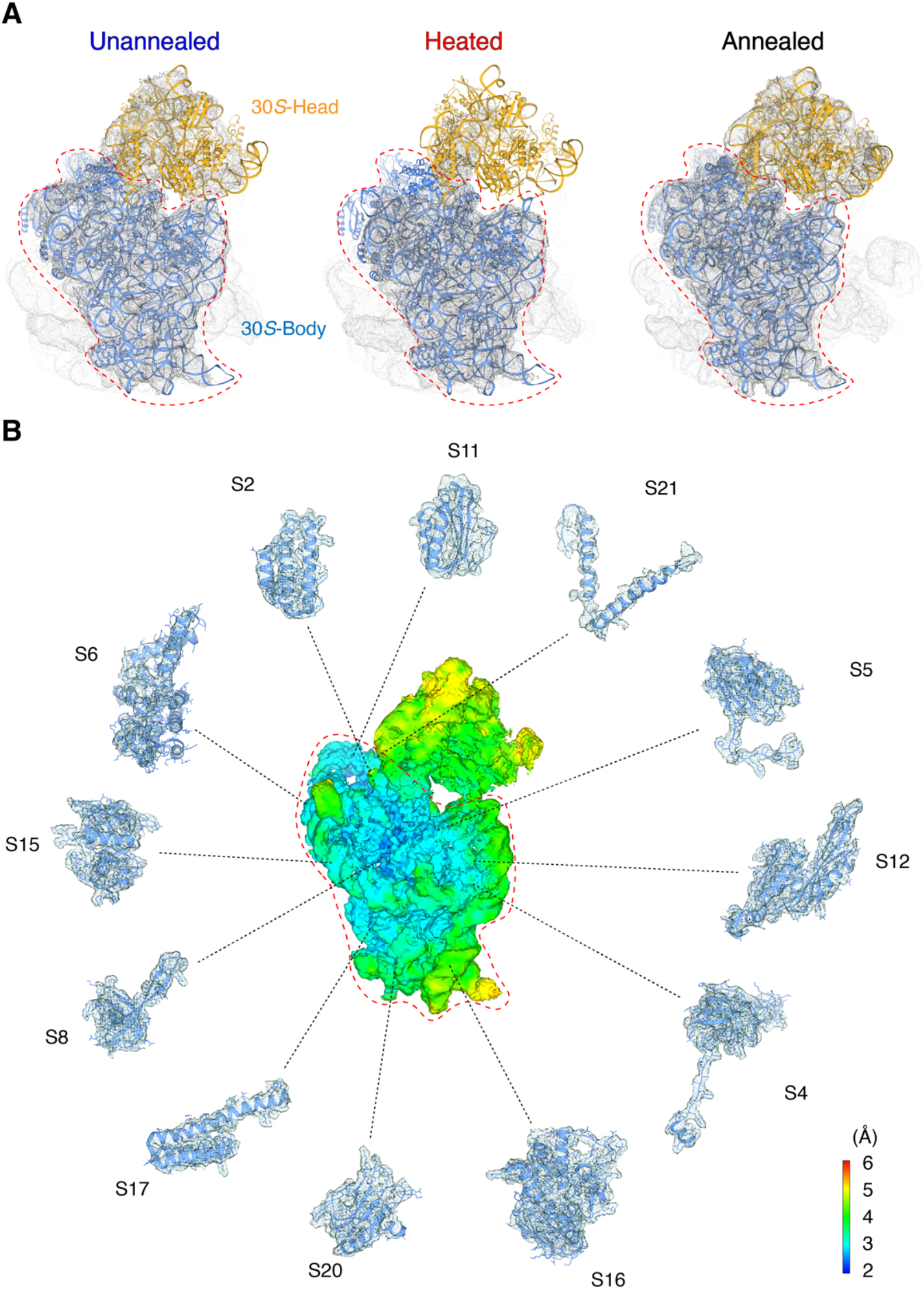
Modeling of 30*S* subunits from the unannealed, heated, and annealed ribosomes. (**A**) Docking of the atomic model of the 30*S* subunit into the unannealed, heated, and annealed ribosomes. The body domains of the 30*S* subunit were marked for high docking accuracy in all ribosomes. (**B**) Expanded view of peripheral proteins fitting into the body domain of the annealed 30*S* subunit.

**Fig. S10.**
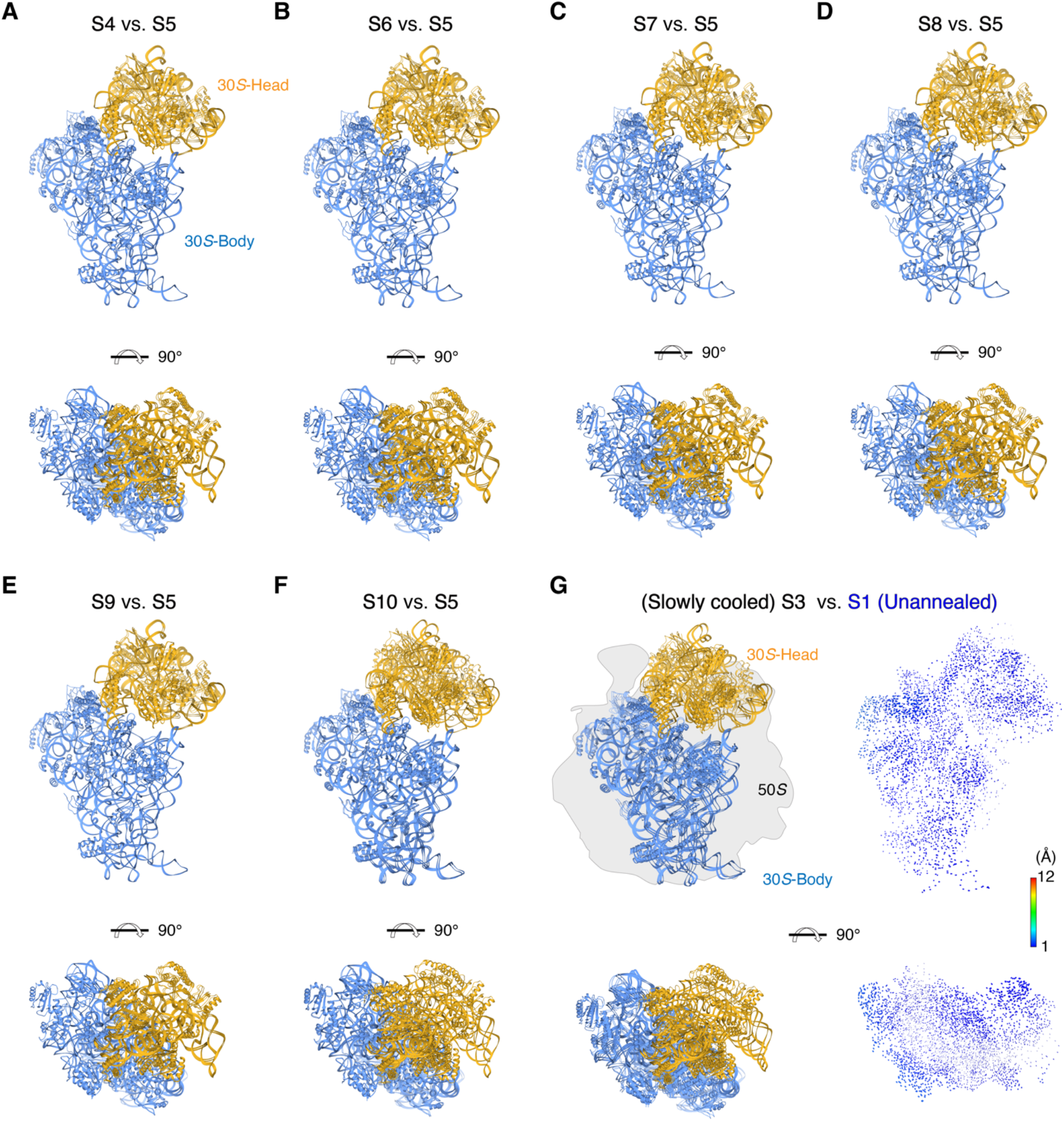
Rotational comparison of 30*S* subunits under various annealing conditions. (**A**–**F**) 50*S* subunits under various annealing conditions were aligned as the reference, and atomic models for the 30*S* subunits are shown. S1, S3, and S4–S10 correspond to annealing conditions as listed in Fig. 2B. (**G**) Rotational comparison of 30*S* subunits between slowly cooled and unannealed ribosomes. Left: Atomic models for the 30*S* subunits. Right: Difference vectors between phosphorous and C*α* atoms in the 30*S* subunits.

**Fig. S11.**
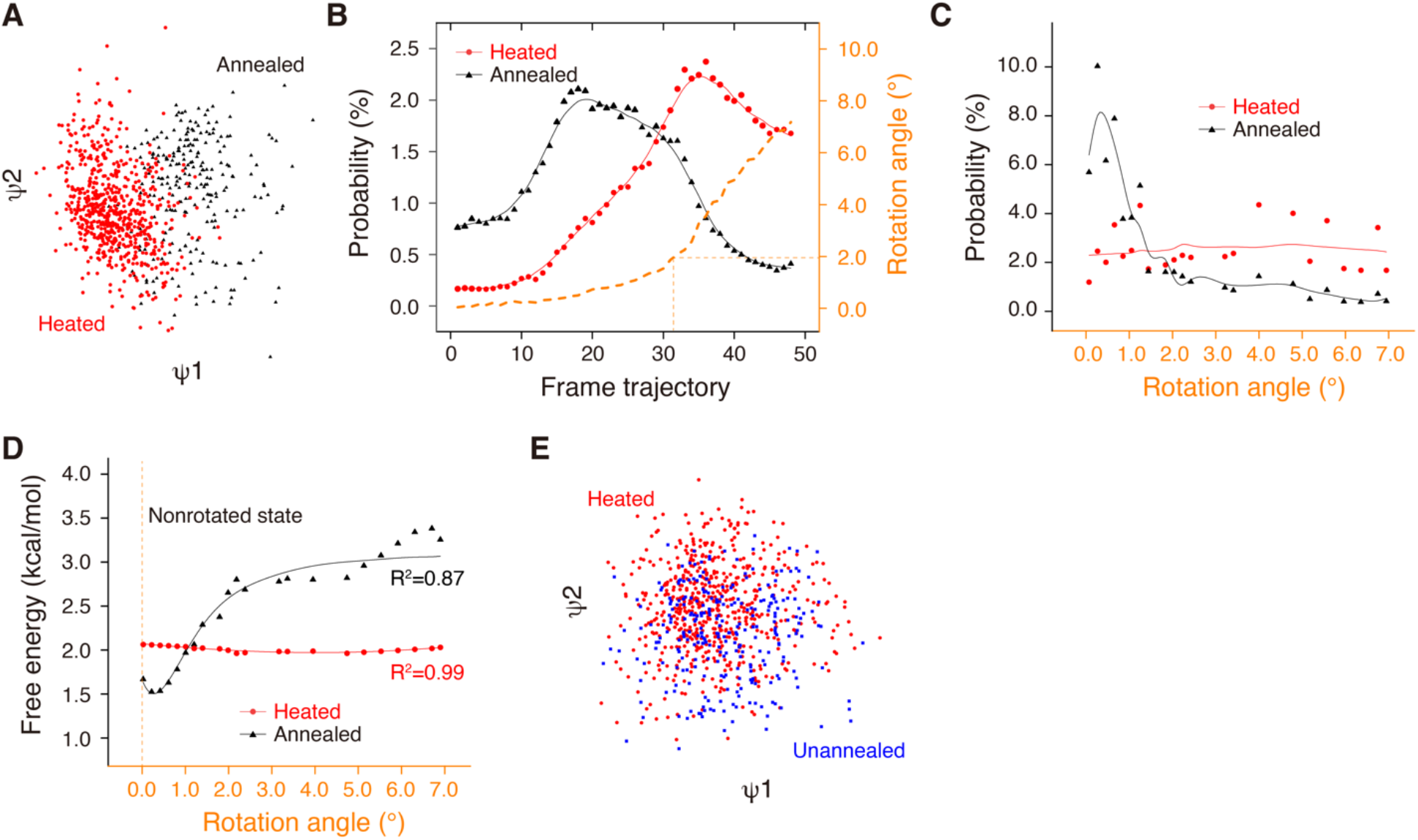
Free-energy minimization occurs during cooling from the heated ribosome to the annealed ribosome. (**A**) Initial manifold snapshots of the 70*S* ribosome in one projection direction. The points are colored in accordance with the heated and annealed subsets. The projection direction is approximately orthogonal to the interface between the 50*S* and 30*S* subunits. (**B**) Particle distribution of the heated and annealed ribosomes along the frame trajectory. The 3D structure at each frame was reconstructed, and the rotation angle of 30*S* subunit with respect to the nonrotated state was calculated. (**C**) Particle distribution of the heated and annealed ribosomes along the rotation angle. Particle distribution was recalculated with the rotation angle at an interval of 0.2°, and a moving average was used to smooth the data variation. (**D**) Free-energy distribution of the heated and annealed ribosomes along the rotation angle. The free energy was calculated from the fitted curve in (**C**). (**E**) Initial manifold snapshots of the 70*S* ribosome in one projection direction. The points are colored in accordance with the unannealed and heated subsets. It is challenging to efficiently separate two states.

**Fig. S12.**
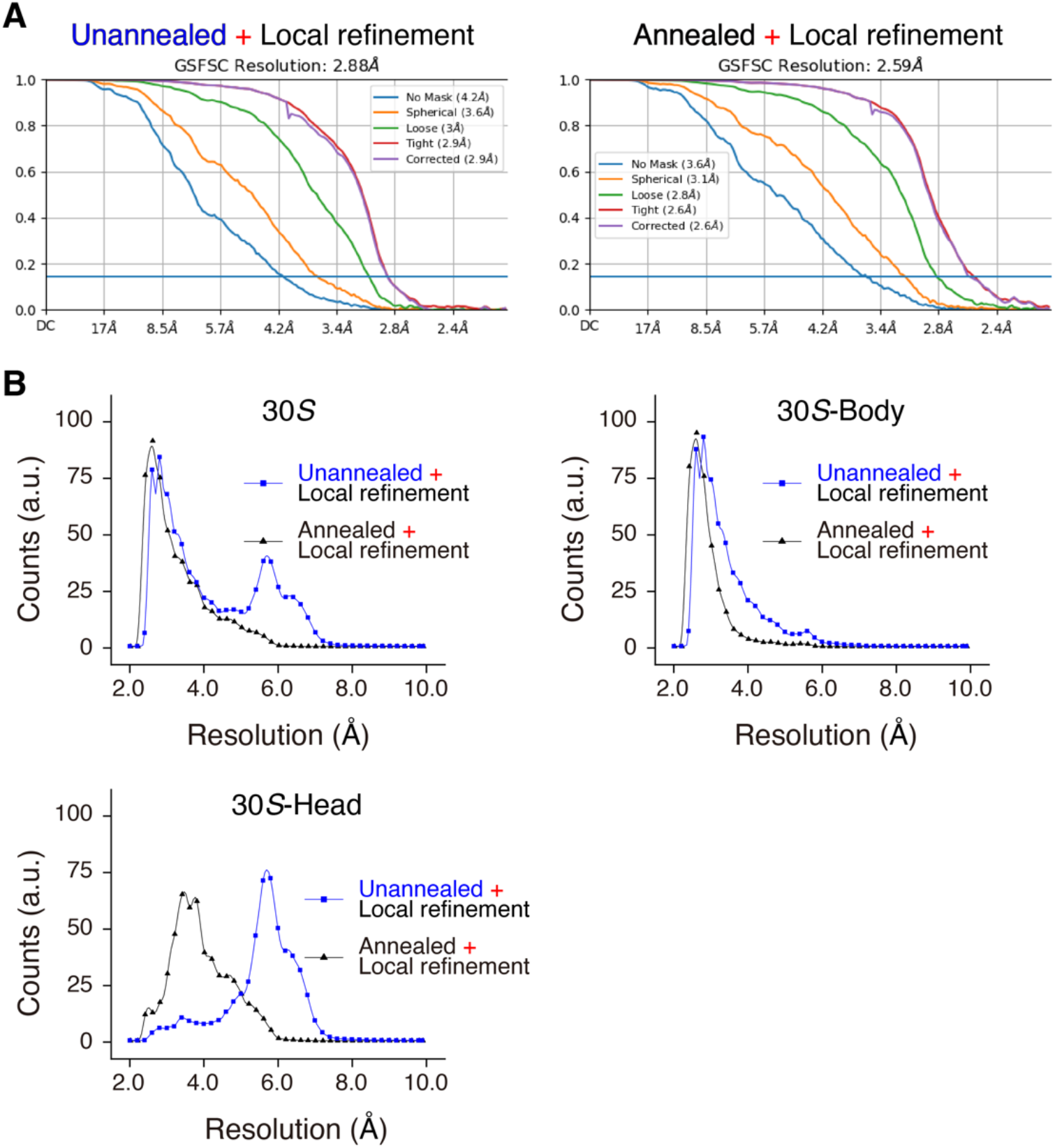
Resolution estimation of 30*S* subunits after local refinements. (**A**) FSC curves for 30*S* subunits after local refinements on unannealed and annealed ribosomes. (**B**) Local resolution histograms of the 30*S* subunit and subdomains after local refinements on unannealed and annealed ribosomes.

**Fig. S13.**
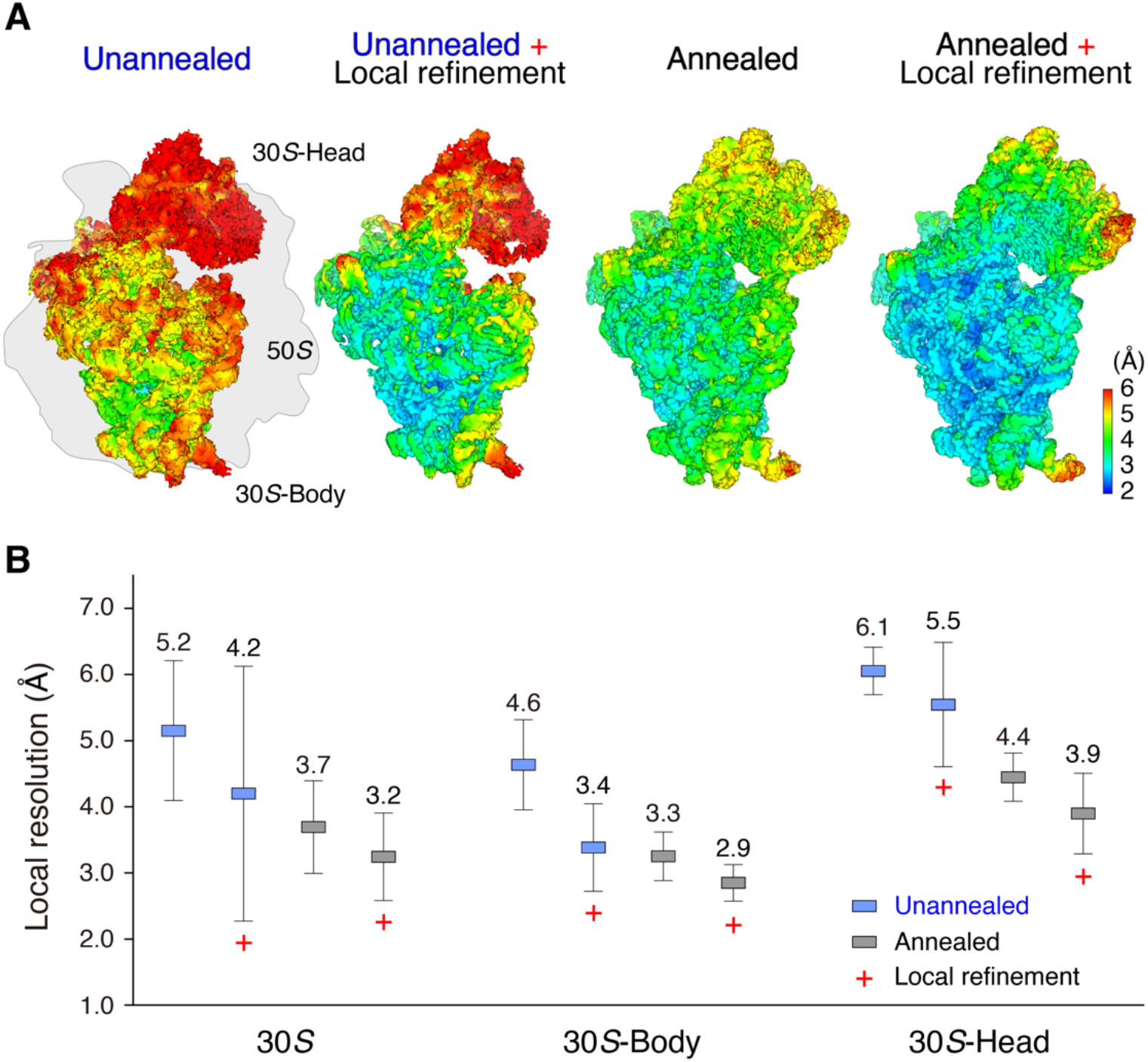
Structural comparison of 30*S* subunits before/after local refinements. (**A**) Local resolution maps of 30*S* subunits under various conditions. (**B**) Local resolution comparison of unannealed and annealed 30*S* subunits before/after local refinements.

**Table S1.**
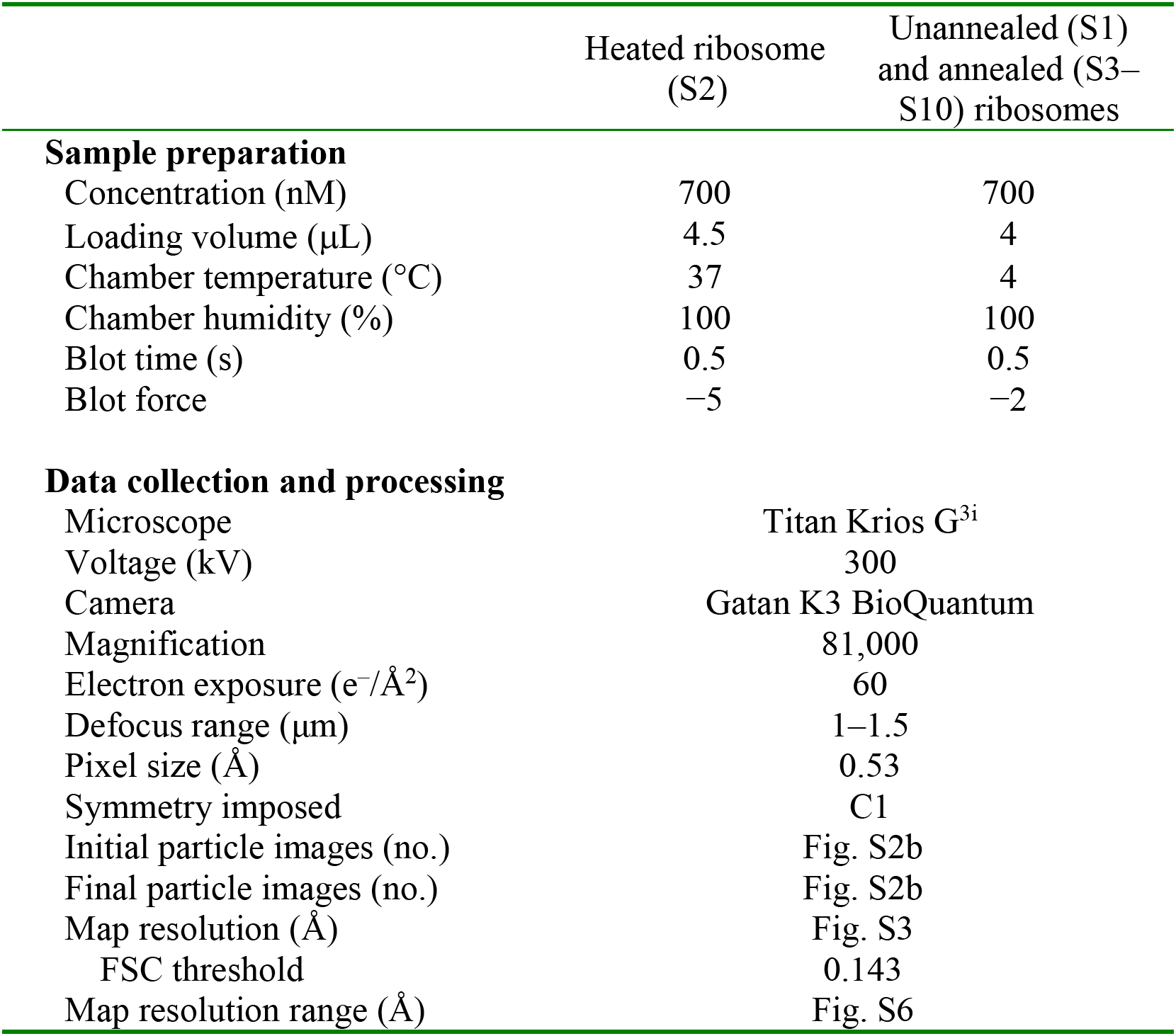
Cryo-EM data collection and data processing statistics.

|  | Heated ribosome<br>(S2) | Unannealed (S1)<br>and annealed (S3–<br>S10) ribosomes |
| --- | --- | --- |
| **Sample preparation** |  |  |
| Concentration (nM) | 700 | 700 |
| Loading volume (μL) | 4.5 | 4 |
| Chamber temperature (°C) | 37 | 4 |
| Chamber humidity (%) | 100 | 100 |
| Blot time (s) | 0.5 | 0.5 |
| Blot force | –5 | –2 |
| **Data collection and processing** |  |  |
| Microscope | Titan Krios G<sup>3i</sup> |  |
| Voltage (kV) | 300 |  |
| Camera | Gatan K3 BioQuantum |  |
| Magnification | 81,000 |  |
| Electron exposure (e<sup>–</sup>/Å<sup>2</sup>) | 60 |  |
| Defocus range (μm) | 1–1.5 |  |
| Pixel size (Å) | 0.53 |  |
| Symmetry imposed | C1 |  |
| Initial particle images (no.) | Fig. S2b |  |
| Final particle images (no.) | Fig. S2b |  |
| Map resolution (Å) | Fig. S3 |  |
| FSC threshold | 0.143 |  |
| Map resolution range (Å) | Fig. S6 |  |

**Movie S1. Cartoon that illustrates how to use the mist umbrella**

**Movie S2. Typical movies for structural variation of the unannealed, heated, and annealed ribosomes**

